# Towards a translationally-independent RNA-based synthetic oscillator using deactivated CRISPR-Cas

**DOI:** 10.1101/2020.05.13.094730

**Authors:** James Kuo, Ruoshi Yuan, Carlos Sánchez, Johan Paulsson, Pamela A. Silver

## Abstract

In synthetic circuits, CRISPR-Cas systems have been used effectively for endpoint changes from an initial state to a final state, such as in logic gates. Here, we use deactivated Cas9 (dCas9) and deactivated Cas12a (dCas12a) to construct dynamic RNA ring oscillators that cycle continuously between states over time in bacterial cells. While our dCas9 circuits using 103-nucleotide guide RNAs showed irregular fluctuations with a wide distribution of peak-to-peak period lengths averaging ∼9 generations, a dCas12a oscillator design with 40-nucleotide CRISPR RNAs performed much better, having a strongly repressed off-state, distinct autocorrelation function peaks, and an average peak-to-peak period length of ∼7.5 generations. Along with free-running oscillator circuits, we measure repression response times in open-loop systems with inducible RNA steps to compare with oscillator period times. We track thousands of cells for 24+ hours at the single-cell level using a microfluidic device. In creating a circuit with nearly translationally-independent behavior, as the RNAs control each others’ transcription, we present the possibility for a synthetic oscillator generalizable across many organisms and readily linkable for transcriptional control.

## Introduction

Oscillators are essential in Nature – circadian rhythm, cell cycles and division, heartbeats, breathing. Human-made oscillators also have a history of great importance, from pendulums in clocks, to crystal oscillators in electronics, to pacemaker and ventilator biomedical devices that interface with the human body. Developing a new biological oscillator may be powerful in many applications, as well as for learning fundamental principles to design synthetic biological circuits with controllable, predictable performance. For example, a synthetic oscillator could be coupled to added functions in enhanced gut bacteria, such as small-molecule drug production (Claesen and Fischbach 2015), lysis for cargo delivery (Din et al. 2016), or timekeeping (Riglar et al. 2019). Protein-based synthetic oscillators have been powerful demonstrations (Stricker et al. 2008, Danino et al. 2010, Potvin-Trottier et al. 2016, Butzin et al. 2017, Scott et al. 2017, Zhang et al. 2017), but building oscillators using nucleic acids has been far less explored (Srinivas et al. 2017). Here, we use CRISPR-Cas systems to develop new circuits whose performance does not depend on activity of varying translated proteins but rather on a varying transcriptional pool of RNAs.

CRISPR-Cas technologies have been used extensively for genome editing and manipulation (Dominguez et al. 2016). For CRISPR-Cas9, the 160 kD Cas9 enzyme forms a complex with a targeting guide RNA, able to bind to specific regions of DNA with base complementarity and cleave. While natural systems typically use two RNAs – a DNA-targeting guide strand and a Cas9 handle strand – these can be combined into a single guide RNA (sgRNA) with both functions. Deactivated Cas9 (dCas9) has the enzymatic active site for DNA cleavage mutated, allowing for gene repression instead of cutting (Bikard et al. 2013, Larson et al. 2013, Qi et al. 2013). Along with using this nuclease-deficient form as a standalone transcriptional repressor, dCas9 has also been fused to transcription factor domains (ex., VP64 or KRAB) to allow for gene activation or further repression (Pickar-Oliver and Gersbach 2019). This strategy of dCas9-controlled transcriptional repression through specific sgRNAs has been used to create logic gates, memory data storage and other synthetic genetic circuits (Nielsen and Voigt 2014, Jusiak et al. 2016, Gander et al. 2017, Sheth et al. 2017). In these cases, the desired readout was an endpoint, a changed expression level after several hours to a final state.

Here, we build on those endpoint circuits to create free-running ring oscillators, where sgRNAs repress each other’s transcription through dCas9 binding of promoters driving sgRNA production. Similarly, we use a deactivated Cas12a (dCas12a) protein along with its targeting CRISPR RNAs in analogous oscillator designs. CRISPR-Cas12a is another bacterial defense system, with the added enzymatic ability of Cas12a to cleave a CRISPR RNA array in addition to its DNA targeting ability (Zetsche et al. 2015). Circuits based on endpoint behavior, such as sensing and logic gates, have also been made with dCas12a (Kempton et al. 2020).

We use a microfluidics device that allows individual tracking of thousands of cells over many generations: *E. coli* cells are confined in trenches with media flowed through, for diffusive feeding that supports exponential growth and washout of newly divided cells (Wang et al. 2010). In addition to oscillators, we make single-cell measurements of dCas9-sgRNA binding repression times, which have mostly been measured at a population level (Qi et al. 2013, Richardson et al. 2016, Jones et al. 2017, Shibata et al. 2017). We compare these response times in inducible RNA cascades with the observed period lengths of the oscillator designs.

Our free-running dCas12a RNA oscillator fluctuates far more regularly than our dCas9-based designs and shows oscillatory behavior, with significant autocorrelation function (ACF) peaks for the population average. Our circuits represent foundational work towards development of an even more regular oscillator using CRISPR-Cas parts and has the potential to be generalizable in a variety of host organisms across kingdoms.

## Results and Discussion

### Design overview: dCas9 RNA oscillator

We designed a ring oscillator using dCas9 and sgRNAs based on the “repressilator” synthetic circuit (Elowitz and Leibler 2000), a three-component ring oscillator with protein repressor parts. At that time, the repressilator was built using well-characterized protein transcription factors (lacI, tetR, phage cI), more recently greatly improved after reducing plasmid noise and removing protein degradation tags (Potvin-Trottier et al. 2016). Similarly, we chose from a characterized set of orthogonal sgRNAs and targeted promoters used in making logic gates with dCas9 repression (Nielsen and Voigt 2014) to assemble a ring oscillator (Fig. 1a). The sgRNAs are expressed from strong bacterial σ^70^ promoters with variations for sgRNA binding between the conserved −35 and −10 regions (Fig. S1), with reported repression fold-changes of 56-250 (Nielsen and Voigt 2014). The protospacer adjacent motif (PAM) sites for dCas9 binding are also between the −35 and −10. Each sgRNA consists of a 20 nucleotide (nt) targeting region and an 83 nt Cas9 handle plus terminator. Additional strong synthetic terminators used previously were also inserted. To preserve any supercoiling effects (Yeung et al. 2017), terminators were set close to the promoter of the downstream sgRNA, separated by several bases from a restriction site. *Streptococcus pyogenes* dCas9 (with deactivating mutations D10A and H840A) was placed under control of an anhydrotetracycline (aTc) inducible TetR promoter (Fig. 1b), as used in other systems (Qi et al. 2013, Nielsen and Voigt 2014).

**Figure 1.**
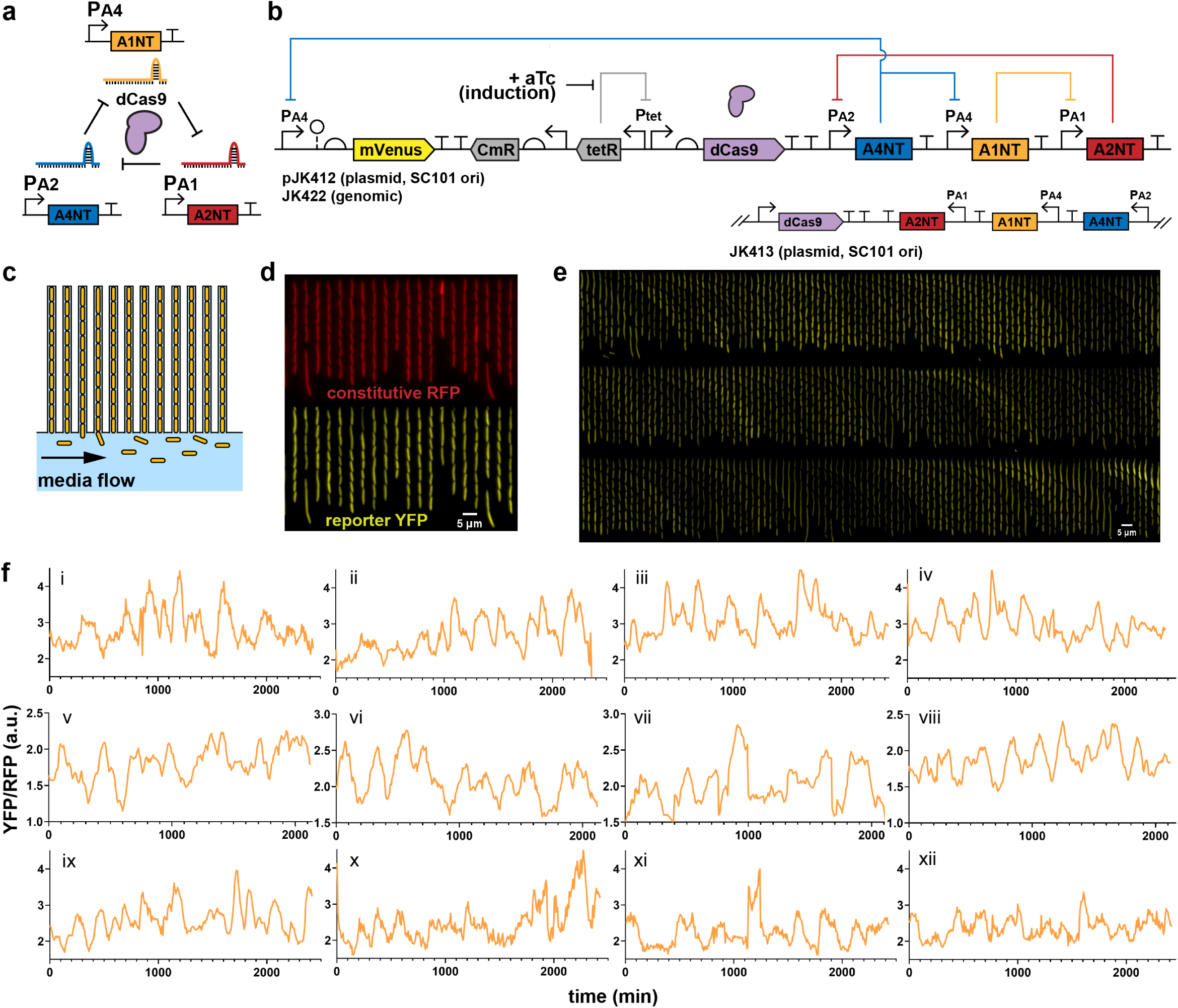
dCas9 RNA oscillator imaged in microfluidic device. (a) The ring oscillator has guide RNAs targeting each other through deactivated Cas9 (dCas9), which forms a complex with a single guide RNA (sgRNA) to create an active transcriptional repressor. Each guide RNA targets the sigma factor binding region of another promoter, which produces its own sgRNA. (b) A design of the dCas9 RNA oscillator with YFP fluorescent reporter mVenus, Tet-inducible dCas9 and the sgRNA ring. The circuit is combined in a single low-copy plasmid (pSC101 origin of replication), genome-integrated, or split across two plasmids. Another form has the sgRNAs transcriptionally reversed. (c) The “mother machine” microfluidic device allows imaging of thousands of cells. Cells are confined in trenches, where “mother” cells are perpetually trapped to allow for single-cell tracking over many generations. (d) *E. coli* MG1655 cells with the designed dCas9 RNA oscillator plasmid (pJK412) are in a mother machine device, with constitutive RFP expressed genomically as a segmentation marker and YFP reporter expressed from the circuit. A field-of-view at one timepoint (6 hours post-aTc induction) shows many cells. (e) In a kymograph looking at a single trench over time, YFP reporter fluorescence appears to fluctuate. Each frame is 8 min, with one generation ∼32 min. (f) Example representative traces of mother cell reporter YFP fluorescence normalized by segmentation RFP signal show fluctuations over time for (i-iv) 1-plasmid circuit (pJK412), (v-viii) genome-integrated circuit (JK422) and (ix-xii) 1-plasmid circuit with flipped sgRNA direction (pJK413).

A yellow fluorescent protein (YFP) mVenus reporter was chosen for its fast maturation rate (Balleza et al. 2018), allowing dynamics to be captured, and placed on the same low-copy plasmid as dCas9. A reporter was made for each of the three different sgRNA promoters, but promoter PA4 outperformed the others in brightness and dynamics, with the lowest reported fold change (Nielsen and Voigt 2014). Different copy number forms of the circuit were created – a two-plasmid system, a one-plasmid version, and a genome-integrated version – with the idea of reducing potential noise from copy number fluctuations.

### Free-running dCas9 RNA oscillator

The dCas9 circuits were transformed into *E. coli* MG1655 with genomic red fluorescent protein (RFP) constitutively expressed as a segmentation marker. Cells were imaged in real time in a “mother machine” microfluidic device that traps cells in trenches while fresh media is flown through, for diffusive feeding that supports exponential growth and washout of newly divided cells (Fig. 1c,d). The mVenus YFP reporter showed fluctuations only with dCas9 induction, where all cells remained bright without added aTc inducer, and these fluctuations occurred continuously for 24+ hours (Fig. 1e; Supplementary Video S1). The RFP segmentation marker did not fluctuate significantly compared to YFP reporter and was used for reporter signal normalization of gene expression noise (Fig. S2). Tracking single mother cells, irregular fluctuations resembling oscillations were observed, with some cells appearing to have some level of regularity (Fig. 1f). A genome-integrated form of the circuit and a form where sgRNAs were transcribed convergent to dCas9 behaved similarly. A two-plasmid system also showed similar fluctuations (Fig. S3), and using a different reporter promoter had much lower signal and dynamic range (Fig. S4). We also tried a different set of orthogonal sgRNAs, which had similar behavior but repressed too strongly for very low reporter signal and range (Fig. S5).

With the wide range of fluctuations across cells, the population-averaged autocorrelation function (ACF) did not show correlation peaks characteristic of oscillators, but the slow decays of multiple generations showed a history-dependence of the reporter signal (Fig. 2). The population-averaged ACF needed over 14 generations to decay to zero. For individual traces that did show a first correlation peak, the average was 8.6 generations. These traces had many different characteristic periods estimated from first correlation peaks (Fig. 2d), which were averaged out when looking over the whole population (Fig. 2c).

**Figure 2.**
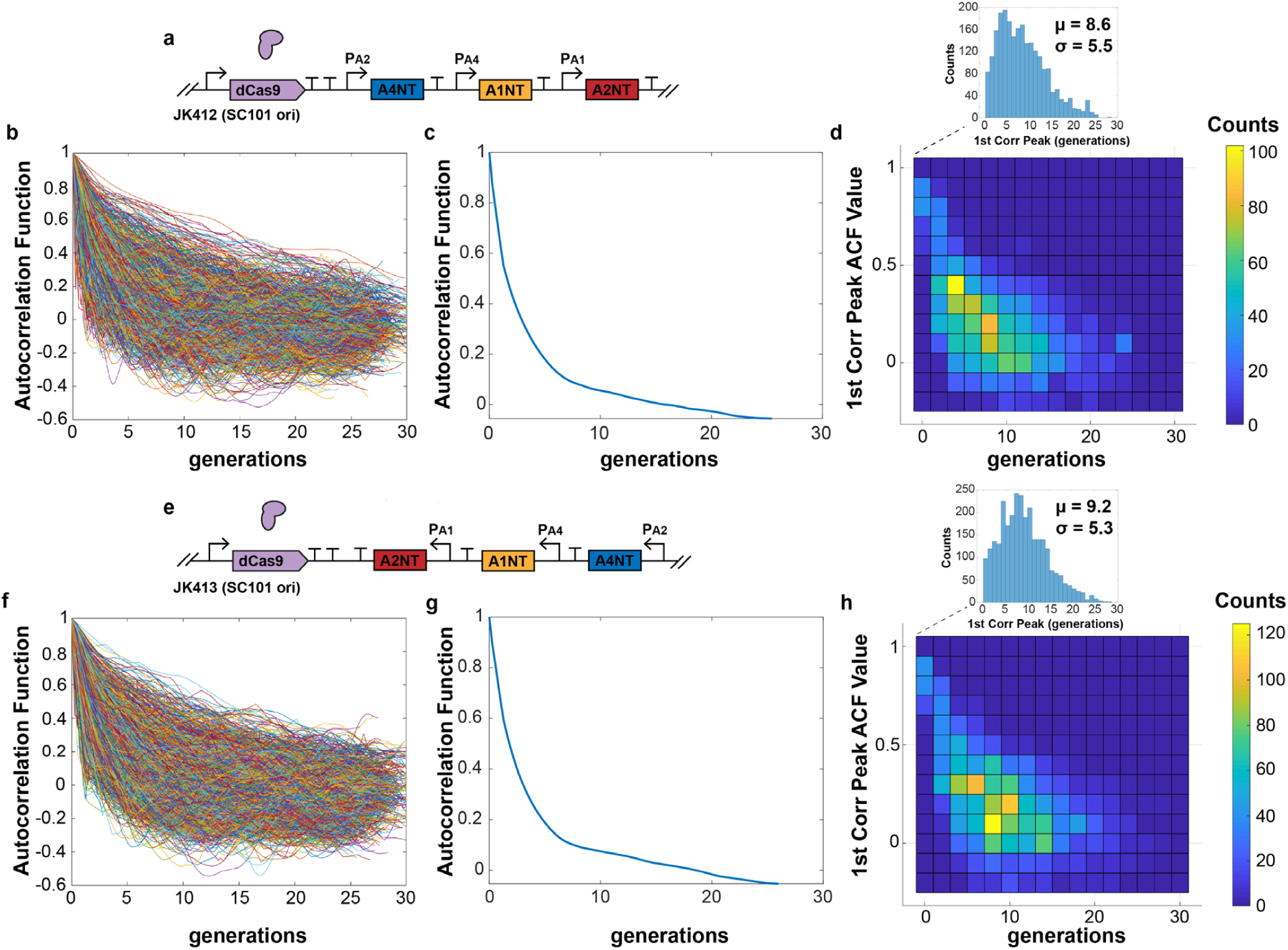
dCas9 RNA oscillator autocorrelation function. (a) The 1-plasmid dCas9 RNA oscillator circuit had autocorrelation function (ACF) calculated for individual cell traces (b). (c) The population-averaged ACF shows a slower decay. (d) Some individual traces showed first correlation peaks across a wide range, with an average of 8.6 generations. (e-h) The same properties were seen for a version with sgRNAs transcriptionally flipped. Cells measured: (a) JK412, 2358; (b) JK413, 3037.

The dCas9-sgRNA complex is fairly stable with nanomolar dissociation constants, predominantly unbinding only during DNA replication (Josephs et al. 2015, Richardson et al. 2016, Jones et al. 2017). This helps explain strong multi-generational repression, as a single dCas9-sgRNA complex could bind to a DNA target for the duration of growth between divisions. Many of the cells did not appear to oscillate obviously, perhaps due to these long binding lifetimes: binding of a lower-population, “wrong” sgRNA could throw off the ring cycle, since the dCas9-sgRNA could effectively bind for an entire generation. This is contrasted with protein repressors such as LacI that individually bind for minutes at a time (Hammar et al. 2014). As with the protein-based repressilator, fluctuations were asynchronous across a population. An effort to synchronize the system by initially producing a high level of one sgRNA from another plasmid failed to entrain the population (Fig. S6).

### Measuring response times for individual dCas9-sgRNA repression steps

To compare with our observed oscillator periods, we created inducible guide RNA circuits using PhlF protein repressor, with an inducible promoter by small molecule 2,4-diacetylphloroglucinol (DAPG). A PhlF promoter replaced an sgRNA promoter in the free-running version for an open-loop version with chemically-inducible control of an sgRNA species (Fig. 3). We could control production of both dCas9 (with aTc) and sgRNA (with DAPG) separately, which we used to characterize response times of individual cells in mother machine experiments for a global single-cell look at dCas9 kinetics.

**Figure 3.**
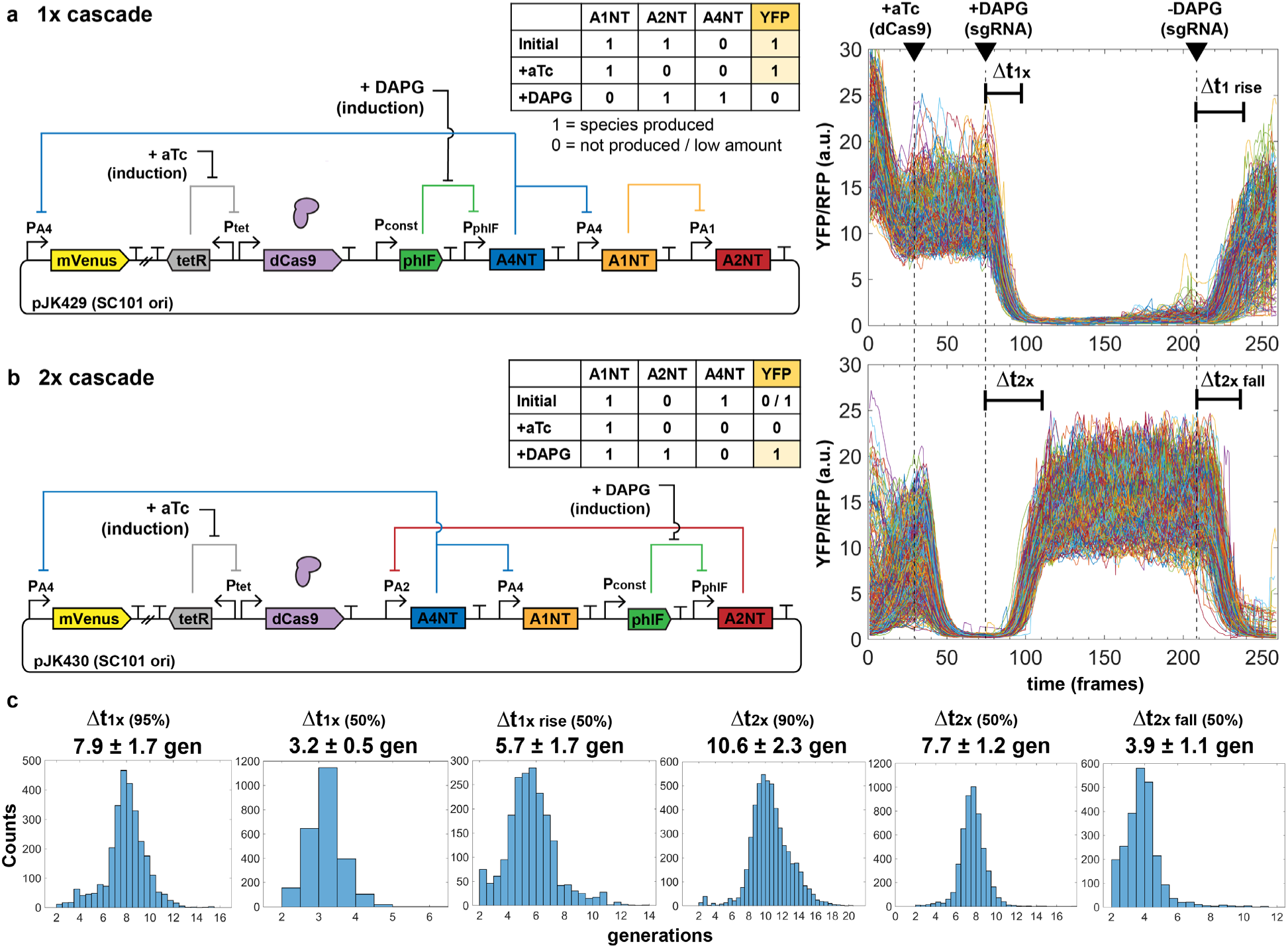
Inducible sgRNA response times are consistent with the free-running oscillator. A single constitutive promoter (PAx) of the free-running system was replaced with a PhlF repressor-based promoter system, allowing inducible control with small molecule DAPG. Measurement of cells in the mother machine allows explicit tracking of individual cell responses to added inducers to determine characteristic times Δt of sgRNA repression cascade steps. *E. coli* MG1655 cells with plasmids were grown in EZ-RDM. (a) One-step sgRNA repression is measured. Expected behavior is YFP bright to dim upon DAPG induction (A4NT sgRNA). Cells are re-entering exponential growth and equilibrating prior to aTc induction. Plot shows 599 cell reporter traces of plasmid version (pJK429). 1 frame = 6 minutes. (b) Two-step sgRNA repression is measured. Expected behavior is YFP dim to bright upon DAPG induction (A2NT sgRNA). Plot shows 515 cell traces of plasmid version (pJK430). (c) Histograms of measured characteristic time steps indicated in (a) and (b). Percentages are percent repression or de-repression. Distributions were measured from (a) 2819 cells for pJK429 and (b) 3136 cells for pJK430.

In our system, we see repression times of several generations dependent on dCas9 expression (Fig. 3). As with the oscillator designs, signal changes were from YFP reporter while constitutive RFP segmentation marker was mostly stable (Fig. S7). Times measured were similar in genome-integrated forms, with lower signal levels (Fig. S8), and in a two-plasmid cascade (Fig. S9). Interestingly, the time difference between the 1x and 2x cascades should be roughly the time for one sgRNA step, as the 1x time includes de-repression of phlF promoter. This time difference is about 2.7 generations (10.6 – 7.9 generations). For the three steps of the free running oscillator, this corresponds to 2.7 × 3 = 8.1 generations, similar to the ∼8.6 generation average period seen for the oscillator plasmid design JK412 (Fig. 2d). Of course, the large variations for the measured oscillator periods make this comparison difficult but suggestive nonetheless.

To our knowledge, this is one of the first instances of tracking dCas9 binding repression in single bacterial cells over time. The times are consistent with those measured by others in different systems at a population level (Qi et al. 2013).

### Exploring dCas9 degradation

With tight dCas9 binding and off rates longer than *E. coli* division times, we thought dCas9 protein degradation might help remove dCas9-sgRNA complexes to more accurately reflect a dynamically changing pool of sgRNAs. We tried adding ssrA (small stable RNA A) degradation tags of varying strengths (Andersen et al. 1998), but did not see improved oscillatory behavior (Fig. S10). Because of this, our main designs use an untagged dCas9. However, roughly setting dCas9 expression level with concentration of aTc inducer was important (Figs. S11-12).

### dCas12a RNA oscillator

CRISPR-Cas12a systems also use a main effector enzyme that acts on DNA by forming a complex with targeting CRISPR RNAs (crRNA) and binding to DNA. We used the ∼150 kD Cas12a protein from *Francisella novicida*, with a nuclease-deactivating D917A mutation (Zetsche et al. 2015, Miao et al. 2019) to create another ring oscillator circuit. The single mutation preserves crRNA array processing ability. Here we use a 19-nt direct repeat as a Cas12a handle along with a 20-nt DNA targeting region (Fig. 4b). The *F. novicida* Cas12a uses a “TTV” protospacer adjacent motif (PAM), which targets the −35 box of the same σ^70^ promoters in the dCas9 designs. We made our Cas12a crRNAs target these promoter regions on the template strand, with the idea that the dCas9 orthogonality with sgRNAs would carry over to the dCas12a system.

**Figure 4.**
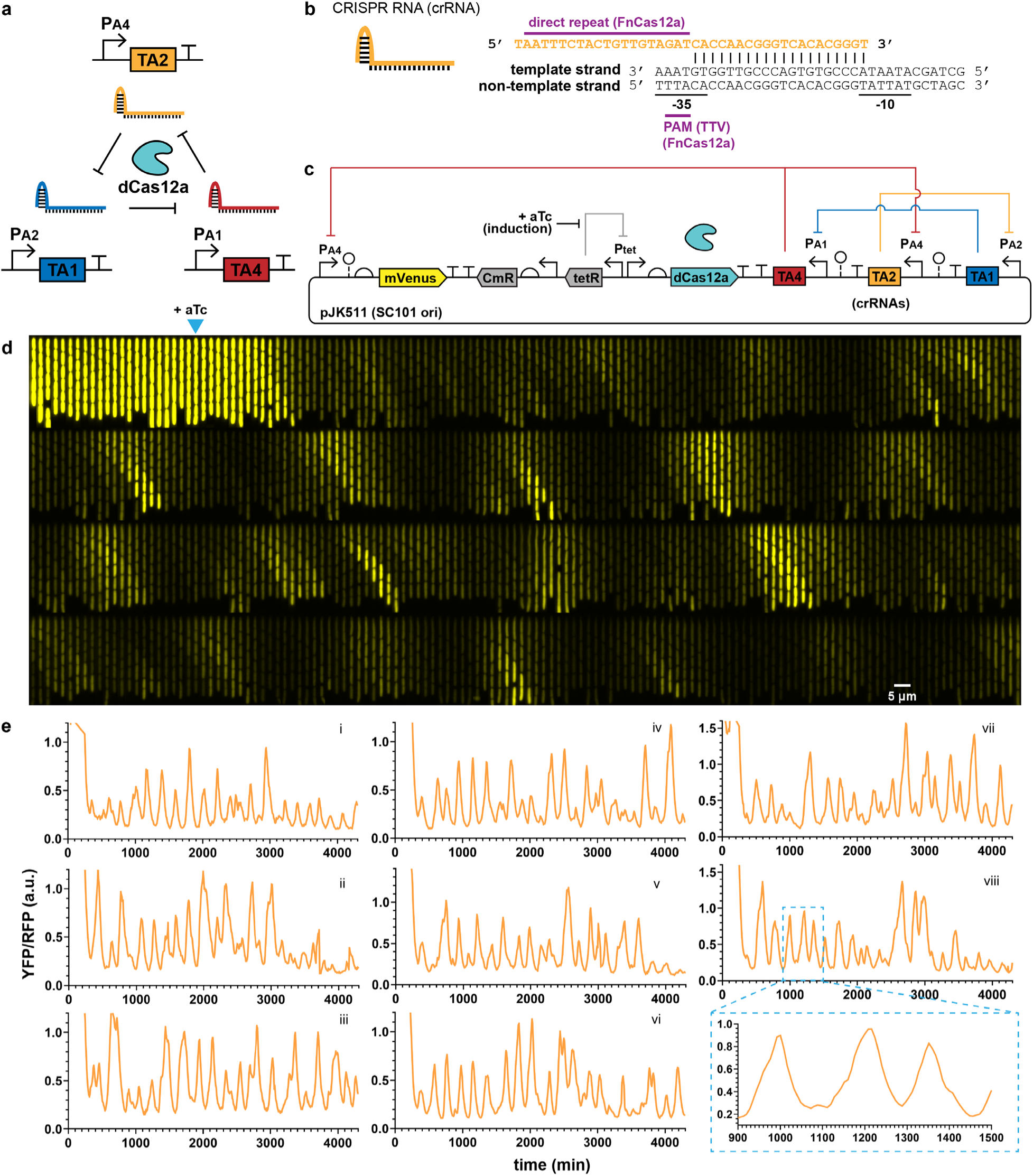
dCas12a RNA oscillator shows regularity. (a) An analogous RNA ring oscillator design replaces dCas9 with deactivated Cas12a (dCas12a). (b) Cas12a CRISPR RNA (crRNA) architecture. Designs use a 20-nt direct repeat dCas12a handle plus 20-nt targeting region. The protospacer adjacent motif (PAM) is “TTV” for *Francisella novicida* Cas12a (FnCas12a). (c) Plasmid design of the dCas12a RNA oscillator circuit has a PA4-mVenus reporter, Tet-inducible dCas12a and crRNAs in a transcriptionally convergent direction. (d,e) *E. coli* MG1655 cells with the circuit were grown in mother machine chips with EZ-RDM. (d) A kymograph of a single trench over time shows YFP reporter fluctuation. Each frame is 8 min, with one generation ∼25 min. Arrow marks induction for dCas12a production at frame 23 (184 min). (e) Example reporter traces show fluctuations resembling oscillations and a lower signal repressed state. Average generation ∼25 min. Pullout for trace viii zooms in on the peak shapes.

As before, we made one- and two- plasmid designs with the same architecture as the dCas9 RNA oscillator (Fig. 4c). To accommodate the shorter 40-nt targeting RNAs for Cas12a, instead of 103-nt for dCas9, we added ∼80-nt ribozyme spacer sequences (Lou et al. 2012) between crRNAs to reduce transcriptional readthrough effects. Notably, strains fluctuated with a much lower baseline, meaning cells looked darker for most of the run with pulses of bright (Fig. 4d,e; Supplementary Video S2). Overall, the fluctuations were much more regular, with some cells appearing quite oscillatory (Fig. 4e). Individual cell ACF traces showed multiple correlation peaks, and the averaged ACF for the population also showed multiple distinct peaks (Fig. 5). We saw a first correlation peak at 7.5 generations and second peak at 14.6 generations. The amplitude of the ACF peaks decays rapidly at first to 0.200, and then drops in smaller increments (∼2.4-fold), similarly to an improved protein repressilator with unstable plasmid removal (Potvin-Trottier et al. 2016). The period length of ∼7.5 generations was similar to the dCas9 oscillators, but slightly faster. Unlike the dCas9 forms, individual cell traces had a much narrower distribution (Fig. 5d), showing the more homogenous behavior across cells. Measuring interpeak distances of apparently oscillating cells, the estimated period length of mean ∼7.5 generations was nearly identical to that determined by ACF (Fig. 5e,f). Peak distances changed as regular multiples of this first, with the variance increasing linearly with increasing interpeak distances by periods (Fig. 5g). This indicated that the period lengths were independent, where the circuit did not show memory effects between periods. For interpeak distance of one period, the standard deviation was 2.9 generations, for an estimated coefficient of variation (CV, standard deviation over mean) of 0.39. The estimated peak heights were seemingly exponentially distributed, with a CV of ∼0.68 (Fig. 5h). Peak heights were also seen to be independent, where consecutive peaks did not show memory/carryover effects in amplitude (Fig. 5i).

**Figure 5.**
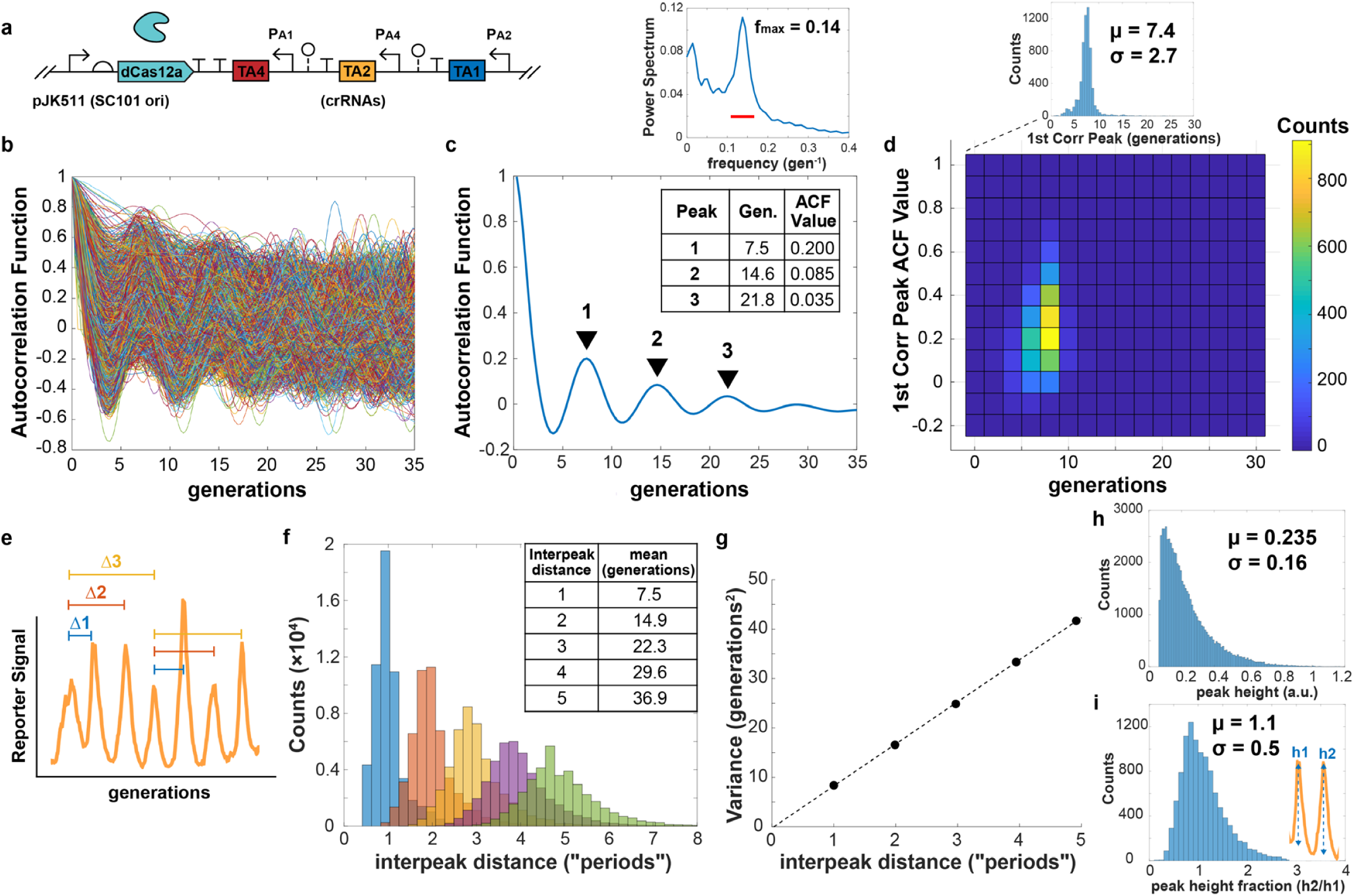
dCas12a RNA oscillator autocorrelation function and interpeak distance. (a) The 1-plasmid dCas12a oscillator circuit with autocorrelation function (ACF) calculated for 6,335 individual cell traces (b). (c) The population-averaged ACF showed clear correlation peaks, with the first correlation peak of ∼7.5 generations. The power spectrum of the ACF shows a max frequency around 0.14, or 7.1 generations. The red bar indicates the width of the window function. (d) Individual cell trace first correlation peaks were mostly clustered around the same period length, with some individual trace ACF peaks showing higher values. (e) Interpeak distance was calculated for peaks of oscillatory traces, where ACF first minimum and first correlation peaks exceeded values ±0.05. (f) Calculated interpeak distances showed distinct distributions following multiples of the first. These are labeled as “periods”, multiples of the 1-interpeak distance. (g) Variances of the interpeak distances increase linearly with period distance. (h) Distribution of estimated peak heights for peaks of oscillatory traces, where ACF first minimum and correlation peaks exceeded ±0.05. (i) Peak height fractions for consecutive peaks, comparing one peak with its preceding one.

Unlike in the dCas9 oscillators, having the crRNAs be transcriptionally convergent with dCas12a had a noticeable improvement in oscillation performance and increased overall brightness, for greater dynamic range of the reporter fluctuations (Fig. S13; Supplementary Video S3). A two-plasmid version of the circuit also showed oscillatory-like fluctuations but was not as regular as the one-plasmid form at the population level (Fig. S14; Supplementary Video S4). Also unlike the dCas9 case, genome-integrating the circuit led to much poorer performance (Fig. S15), suggesting the higher copy numbers of the plasmid are needed for the dCas12a oscillatory behavior.

### Measuring response times for individual dCas12a-crRNA repression steps

As with the dCas9 system, we created inducible crRNA circuits using PhlF protein repressor, to compare with our measured periods of the observed fluctuations. Repression times were characterized as with the dCas9 systems, with a one-step repression time of ∼9 generations (Fig. 6). However, the time for 50% repression was rapid at only ∼2 generations, likely accounting for the faster cycling time of the oscillator. As with dCas9, dCas12a-crRNA repression of YFP began soon after crRNA induction, and the repression was strong at ∼36-fold (1.5 to 0.41 a.u. signal change high to low) and maintained in most cells for at least 600 minutes (∼25 generations).

**Figure 6.**
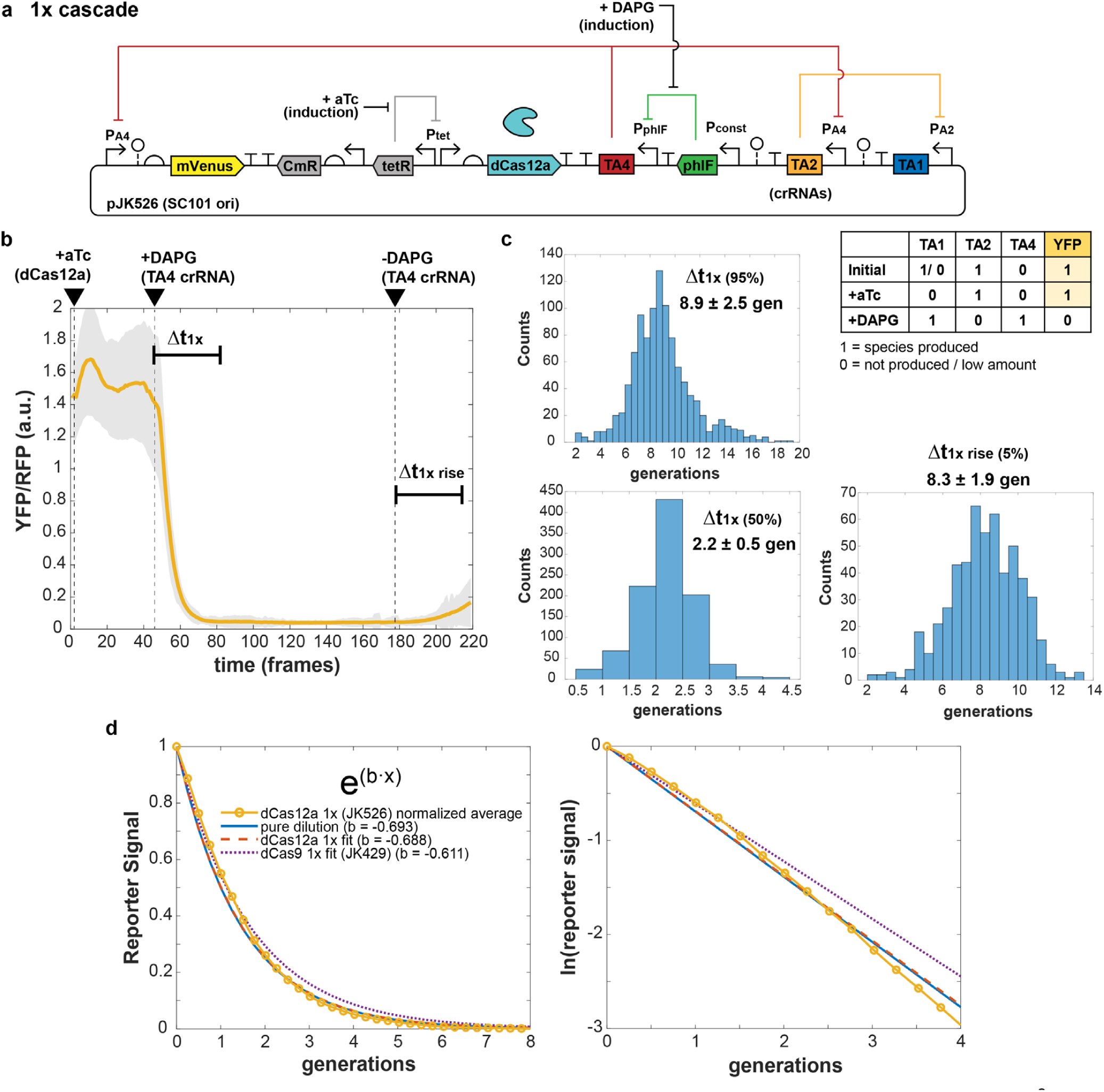
dCas12a crRNA repression response times. (a) A single constitutive promoter of the free-running dCas12a oscillator was replaced with a PhlF repressor-based promoter system, allowing inducible control with small molecule DAPG and measurement of characteristic times Δt of crRNA repression cascade steps. *E*.*coli* MG1655 cells with plasmids were grown in EZ-RDM in the mother machine. (b) One-step crRNA repression is measured. Expected behavior is YFP bright to dim upon DAPG induction (TA4 crRNA). (b) Averaged trace from 1049 cells of plasmid version (pJK526), with shaded region showing one standard deviation of the averaged values. (Single traces in Fig. S16.) 1 frame = 6 minutes. (c) Histograms of measured characteristic time steps indicated in (b) for single cell traces. Percentages are percent repression or de-repression. Distributions measured from 1045 cells. (d) Exponential fitting of the averaged trace (JK526) and dCas9 1x cascade trace. The “pure dilution” curve (solid blue) corresponds to that expected for dilution from cell division alone.

Fitting the average trace with a simple exponential (a·exp(b·t)), the exponent coefficient of b = −0.688, was very close to that expected for pure dilution based on cell division alone (b = - ln(2) = −0.693) (Fig. 6d). This was also close to that observed from the averaged trace of the dCas9 single step cascade (b = −0.611). These results show that in these inducible step strains, the reporter decay processes are nearly pure dilution from cell division.

A two-step crRNA cascade circuit did not behave as expected (Fig. S17). Cell reporter signal was repressed upon aTc addition for dCas12a induction as expected, but unexpectedly then steadily increased before crRNA induction. The steady-state signal with crRNA induction was about half that of before dCas12a induction, and removal of crRNA inducer caused only a small drop in signal. Cells grew normally, with average division time of ∼25.8 minutes. The unexpected output suggested some feedback coupling between the crRNAs, likely from transcriptional readthrough in spite of the strong terminators and ribozyme spacer sequences. Some level of this crRNA coupling is likely also present in the oscillator, though unclear how it affects performance. For the dCas9 circuits, an extra transcriptional terminator (40 nt, from *S. pyogenes*) is built into each sgRNA by design (Qi et al. 2013) that likely reduces transcriptional readthrough effects.

### Cell growth comparisons

Cell growth rates of dCas12a strains were fairly normal in mother machine runs, with average division times of ∼25.7 minutes for the dCas12a oscillator (Fig. 7a). These rates are close to those without circuit or with repressilator or dual-feedback oscillator circuits of ∼24-25 minutes (Luro et al. 2020), suggesting low circuit burden. In contrast, the dCas9 oscillator strains had slowed growth, with increased division rates up to 30+ minutes. (It was difficult to assess exact growth impacts of individual circuits because other variable factors of media condition and PDMS chip could affect growth rate much more. We had runs where the media and chip conditions increased division rates by a few minutes without changing the strain’s circuit performance by generations timescale.) While division times could vary due to conditions, the dCas12a and dCas9 oscillator strains were from the same mother machine run for direct comparison in the results presented. The oscillator circuits appeared to be greater burden than the inducible RNA circuits, whose strains all grew faster on average. The distributions give a look at the heterogeneity of the populations, with a relatively narrow distribution for the dCas12a oscillator and wide distributions for other strains (Fig. 7b).

**Figure 7.**
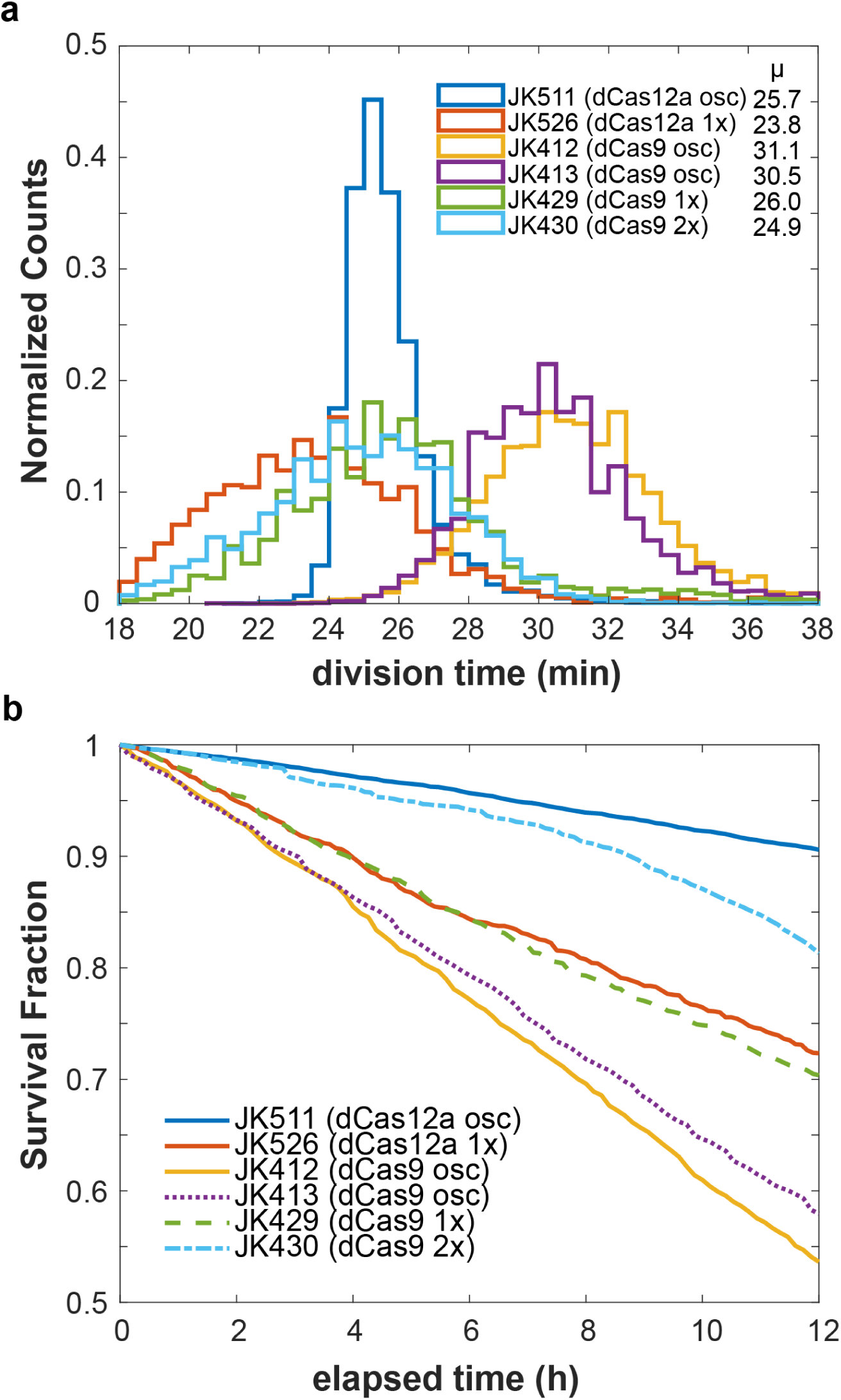
Cell growth comparisons. Estimates on growth efficiency of RNA oscillator and cascade strains. Cell division times depended on media and chip conditions. (a) Average growth division times, as observed in mother machine runs. Each count is a single mother cell’s average division time over the course of the run. Cell counts: JK511, 7998; JK526, 2263; JK412, 7158; JK413, 7396; JK429, 5921; JK430, 7779. (b) “Survival” curves of mother cells, estimated in a 12-hour window within the first 24 hours of imaging. The fraction of mother cell trajectories remaining after the indicated time estimates cell survival.

However, we did see much greater rates of filamentation and cell death from all circuits. By looking at a 12-hour window in the first 24 hours of imaging, we could get an estimate of circuit toxicity based on surviving mother cells. While the dCas12a oscillator (JK511) had over 90% survival, the dCas9 oscillators (JK412, JK413) had only ∼55-60% survival. A circuit without toxicity is expected to be over 99.9% (Luro et al. 2020). As with growth rates, death rates also varied due to chip conditions. The higher rates of cell death, often by filamentation, were likely due to toxicity from one or more of the guide RNAs as dCas9-sgRNA complex (Cho et al. 2018, Cui et al. 2018) and a similar toxicity from dCas12a-crRNA complex. The greater toxicity seen from dCas9 compared to dCas12a is consistent with that seen in other bacteria (Jiang et al. 2017, Knoot et al. 2020). Perhaps toxicity may be reduced through protein engineering efforts, as was done for dCas9 (Zhang and Voigt 2018). It is worth noting that even with these toxicity rates, a growing population will still increase substantially: these death rates are in a 12-h window of 24+ new generations of exponentially growing cells.

## Conclusion

We present construction of new translationally-independent synthetic oscillators based on transcription of CRISPR-associated RNAs. In this manner, we present one of the first dynamic uses of CRISPR-Cas systems over time. The only other example we are aware of is an experimental effort to make a dCas9-based CRISPR interference oscillator (Santos-Moreno et al. 2020), where their periods were much longer at 14-17 generations, as well as a modeling-only exploration (Clamons and Murray 2017). Most previous CRISPR component circuits, while impressive, used endpoint as the behavior, showing before-and-after changes from incubations of several hours or more. That is excellent for sensors and logic gates, but we show a time-varying system that fluctuates between on/off reporter states in an analog manner. In addition, our characterization method is one of the first examples of single-cell analysis of a dCas9 or dCas12a circuit in bacteria (Jones et al. 2017, Martens et al. 2019, Camsund et al. 2020). We also use our setup to explicitly measure single-cell dCas9-sgRNA and dCas12a-crRNA repression times in thousands of individual cells.

While we do not see performance as regular as the high-precision improved protein-based repressilator (Potvin-Trottier et al. 2016), we see clear signs of regularity that are promising for further improvement or immediate applications with less stringent timing requirements. For the dCas12a RNA oscillator, some cells have quite regular timing, and the circuit performs very similarly to an improved version of the protein-based repressilator on a single plasmid, with an even better population-averaged ACF (Potvin-Trottier et al. 2016). This proof-of-concept may allow construction of chained RNA ring oscillators, such as two three-component rings or larger 5+ component rings. Logic gate type functional output can be created with two rings, such as AND or NOR gates where each ring partly controls repression of a single promoter output. We point out that many original proof-of-concept oscillator designs (Elowitz and Leibler 2000, Stricker et al. 2008) had performed irregularly at first but were greatly improved in separate major efforts (Potvin-Trottier et al. 2016, Luro et al. 2020).

The dCas12a RNA oscillator may also be an interesting model system for further study in itself, as the oscillatory behavior is seen without direct binding cooperativity in the circuit. However, the system likely has nonlinearity from other mechanisms, such as bound-repressor degradation (Loinger and Biham 2007), with DNA-bound dCas12a-crRNA removed from cell divisions and possibly protein degradation, or Michaelis-Menten degradation of the RNAs (Ananthasubramaniam and Herzel 2014). Further investigation of these alternatives to cooperative binding for nonlinearity may lead to better understanding and new strategies for designing other oscillator circuits.

In the dCas12a design, another crRNA can be added at the end of any existing oscillator RNA to form a crRNA array. These added crRNAs can target other desired sequences for regulation beyond the core oscillator crRNAs, with the CRISPR array being processed by dCas12a. In this way, our circuits may be readily coupled for transcription regulation controlled by the oscillator, including potential control of unmodified endogenous genes. Other versions of dCas12a may be useful: *Acidaminococcus* sp. BV3L6 (AsCas12a) and *Lachnospiracaeae* bacterium ND2006 (LbCas12a) Cas12a both have more specific PAM targeting sites of “TTTV” (Zetsche et al. 2015). The targeted promoters in our circuits already have a “TTTG” PAM-motif in the −35 region of the strong σ^70^ promoters, so these other Cas12as may reduce any off-target effects. Variants of Cas12a can also target different PAM sites, further increasing the versatility to target many sequences (Kleinstiver et al. 2019, Toth et al. 2020). Also, AsCas12a and LbCas12a have had faster association on-rates measured (Singh et al. 2018), so these may be quick ways to tune the period lengths of the oscillator. The FnCas12a on-rate (k_on_ ∼ 2.3 × 10^6^ M^-1^ ·s^-1^, (Singh et al. 2018)) was measured as roughly half or a third that of Cas9 (k_on_ ∼ 6 × 10^6^ M^-1^·s^-1^, (Singh et al. 2016)), maybe a factor in the dCas12a oscillator’s better performance than dCas9. Addition of ribozyme sequences between the crRNAs, not added for the sgRNA, may have also helped, regulating RNA amounts by insulating from context effects such as from transcriptional readthrough (Nielsen et al. 2016).

We have built prototype oscillators that are nearly translationally-independent in their output behavior. Since dCas9 and dCas12a can be functionally expressed in many unique hosts across kingdoms, a generalizable oscillator circuit may be possible.

## Methods

For detailed descriptions, see Supporting Information.

### Cloning and strain preparation

For the dCas9 system, components were amplified from existing plasmids (Qi et al. 2013, Nielsen and Voigt 2014). For dCas12a, Cas12a pieces were PCR amplified with new D917A mutation from existing plasmid (Zetsche et al. 2015). crRNA components were synthesized (IDT) as <1 kb fragments and combined with Gibson assembly. All pieces were combined with Gibson assembly or restriction enzymes with T4 ligase in *E. coli* DH5a or EC100D pir+ for genomic integrations using pOSIP system (St-Pierre et al. 2013).

### Microfluidic imaging

Mother machine microfluidic chips were cast with PDMS using wafer designs for *E. coli* cells. Most runs used chips with trench dimensions of 1.5 µm wide and high, with different lengths (10+ µm) used throughout. EZ Rich Defined Medium (EZ-RDM, TekNova) was used with 0.85 g/L Pluronic F108 (Sigma-Aldrich) to prevent cell adhesion to PDMS. Media was flown at 20 µL/min through Tygon microfluidic tubing with NE-300 syringe pumps (New Era Pump Systems). Cells were imaged on modified Nikon Ti2-E inverted microscopes with a 40x air objective.

## Acknowledgements

We thank Somenath Bakshi, Emanuele Leoncini, Luis Gutiérrez-López, Laurent Potvin-Trottier, Scott Luro, Jeffrey Way, and other lab members for their help. Some microscopy experiments were performed at The Nikon Imaging Center at Harvard Medical School with help from Jennifer Waters and Anna Payne-Tobin Jost. Thanks also to K.B. This research was supported by Defense Advanced Research Projects Agency (DARPA) Grant HR0011-16-2-0049 and the National Institute of General Medical Sciences of the NIH under a Ruth L. Kirschstein National Research Service Award (F32GM125108 to J.K.).

## References

Ananthasubramaniam, B. and H. Herzel (2014). “Positive feedback promotes oscillations in negative feedback loops.” PLoS One 9(8): e104761.

Andersen, J. B., C. Sternberg, L. K. Poulsen, S. P. Bjorn, M. Givskov and S. Molin (1998). “New unstable variants of green fluorescent protein for studies of transient gene expression in bacteria.” Appl Environ Microbiol 64(6): 2240–2246.

Balleza, E., J. M. Kim and P. Cluzel (2018). “Systematic characterization of maturation time of fluorescent proteins in living cells.” Nat Methods 15(1): 47–51.

Bikard, D., W. Jiang, P. Samai, A. Hochschild, F. Zhang and L. A. Marraffini (2013). “Programmable repression and activation of bacterial gene expression using an engineered CRISPR-Cas system.” Nucleic Acids Res 41(15): 7429–7437.

Butzin, N. C., P. Hochendoner, C. T. Ogle and W. H. Mather (2017). “Entrainment of a Bacterial Synthetic Gene Oscillator through Proteolytic Queueing.” ACS Synth Biol 6(3): 455–462.

Camsund, D., M. J. Lawson, J. Larsson, D. Jones, S. Zikrin, D. Fange and J. Elf (2020). “Time-resolved imaging-based CRISPRi screening.” Nat Methods 17(1): 86–92.

Cho, S., D. Choe, E. Lee, S. C. Kim, B. Palsson and B. K. Cho (2018). “High-Level dCas9 Expression Induces Abnormal Cell Morphology in Escherichia coli.” ACS Synth Biol 7(4): 1085–1094.

Claesen, J. and M. A. Fischbach (2015). “Synthetic microbes as drug delivery systems.” ACS Synth Biol 4(4): 358–364.

Clamons, S. E. and R. M. Murray (2017). “Modeling Dynamic Transcriptional Circuits with CRISPRi.” bioRxiv: 225318.

Cui, L., A. Vigouroux, F. Rousset, H. Varet, V. Khanna and D. Bikard (2018). “A CRISPRi screen in E. coli reveals sequence-specific toxicity of dCas9.” Nat Commun 9(1): 1912.

Danino, T., O. Mondragon-Palomino, L. Tsimring and J. Hasty (2010). “A synchronized quorum of genetic clocks.” Nature 463(7279): 326–330.

Din, M. O., T. Danino, A. Prindle, M. Skalak, J. Selimkhanov, K. Allen, E. Julio, E. Atolia, L. S. Tsimring, S. N. Bhatia and J. Hasty (2016). “Synchronized cycles of bacterial lysis for in vivo delivery.” Nature 536(7614): 81–85.

Dominguez, A. A., W. A. Lim and L. S. Qi (2016). “Beyond editing: repurposing CRISPR-Cas9 for precision genome regulation and interrogation.” Nat Rev Mol Cell Biol 17(1): 5–15.

Elowitz, M. B. and S. Leibler (2000). “A synthetic oscillatory network of transcriptional regulators.” Nature 403(6767): 335–338.

Gander, M. W., J. D. Vrana, W. E. Voje, J. M. Carothers and E. Klavins (2017). “Digital logic circuits in yeast with CRISPR-dCas9 NOR gates.” Nat Commun 8: 15459.

Hammar, P., M. Wallden, D. Fange, F. Persson, O. Baltekin, G. Ullman, P. Leroy and J. Elf (2014). “Direct measurement of transcription factor dissociation excludes a simple operator occupancy model for gene regulation.” Nat Genet 46(4): 405–408.

Jiang, Y., F. Qian, J. Yang, Y. Liu, F. Dong, C. Xu, B. Sun, B. Chen, X. Xu, Y. Li, R. Wang and S. Yang (2017). “CRISPR-Cpf1 assisted genome editing of Corynebacterium glutamicum.” Nat Commun 8: 15179.

Jones, D. L., P. Leroy, C. Unoson, D. Fange, V. Curic, M. J. Lawson and J. Elf (2017). “Kinetics of dCas9 target search in Escherichia coli.” Science 357(6358): 1420–1424.

Josephs, E. A., D. D. Kocak, C. J. Fitzgibbon, J. McMenemy, C. A. Gersbach and P. E. Marszalek (2015). “Structure and specificity of the RNA-guided endonuclease Cas9 during DNA interrogation, target binding and cleavage.” Nucleic Acids Res 43(18): 8924–8941.

Jusiak, B., S. Cleto, P. Perez-Pinera and T. K. Lu (2016). “Engineering Synthetic Gene Circuits in Living Cells with CRISPR Technology.” Trends Biotechnol 34(7): 535–547.

Kempton, H. R., L. E. Goudy, K. S. Love and L. S. Qi (2020). “Multiple Input Sensing and Signal Integration Using a Split Cas12a System.” Mol Cell 78(1): 184–191 e183.

Kleinstiver, B. P., A. A. Sousa, R. T. Walton, Y. E. Tak, J. Y. Hsu, K. Clement, M. M. Welch, J. E. Horng, J. Malagon-Lopez, I. Scarfo, M. V. Maus, L. Pinello, M. J. Aryee and J. K. Joung (2019). “Engineered CRISPR-Cas12a variants with increased activities and improved targeting ranges for gene, epigenetic and base editing.” Nat Biotechnol 37(3): 276–282.

Knoot, C. J., S. Biswas and H. B. Pakrasi (2020). “Tunable Repression of Key Photosynthetic Processes Using Cas12a CRISPR Interference in the Fast-Growing Cyanobacterium Synechococcus sp. UTEX 2973.” ACS Synth Biol 9(1): 132–143.

Larson, M. H., L. A. Gilbert, X. Wang, W. A. Lim, J. S. Weissman and L. S. Qi (2013). “CRISPR interference (CRISPRi) for sequence-specific control of gene expression.” Nat Protoc 8(11): 2180–2196.

Loinger, A. and O. Biham (2007). “Stochastic simulations of the repressilator circuit.” Phys Rev E Stat Nonlin Soft Matter Phys 76(5 Pt 1): 051917.

Lou, C., B. Stanton, Y. J. Chen, B. Munsky and C. A. Voigt (2012). “Ribozyme-based insulator parts buffer synthetic circuits from genetic context.” Nat Biotechnol 30(11): 1137–1142.

Luro, S., L. Potvin-Trottier, B. Okumus and J. Paulsson (2020). “Isolating live cells after high-throughput, long-term, time-lapse microscopy.” Nat Methods 17(1): 93–100.

Martens, K. J. A., S. P. B. van Beljouw, S. van der Els, J. N. A. Vink, S. Baas, G. A. Vogelaar, S. J. J. Brouns, P. van Baarlen, M. Kleerebezem and J. Hohlbein (2019). “Visualisation of dCas9 target search in vivo using an open-microscopy framework.” Nat Commun 10(1): 3552.

Miao, C., H. Zhao, L. Qian and C. Lou (2019). “Systematically investigating the key features of the DNase deactivated Cpf1 for tunable transcription regulation in prokaryotic cells.” Synth Syst Biotechnol 4(1): 1–9.

Nielsen, A. A., B. S. Der, J. Shin, P. Vaidyanathan, V. Paralanov, E. A. Strychalski, D. Ross, D. Densmore and C. A. Voigt (2016). “Genetic circuit design automation.” Science 352(6281): aac7341.

Nielsen, A. A. and C. A. Voigt (2014). “Multi-input CRISPR/Cas genetic circuits that interface host regulatory networks.” Mol Syst Biol 10: 763.

Pickar-Oliver, A. and C. A. Gersbach (2019). “The next generation of CRISPR-Cas technologies and applications.” Nat Rev Mol Cell Biol 20(8): 490–507.

Potvin-Trottier, L., N. D. Lord, G. Vinnicombe and J. Paulsson (2016). “Synchronous long-term oscillations in a synthetic gene circuit.” Nature 538(7626): 514–517.

Qi, L. S., M. H. Larson, L. A. Gilbert, J. A. Doudna, J. S. Weissman, A. P. Arkin and W. A. Lim (2013). “Repurposing CRISPR as an RNA-guided platform for sequence-specific control of gene expression.” Cell 152(5): 1173–1183.

Richardson, C. D., G. J. Ray, M. A. DeWitt, G. L. Curie and J. E. Corn (2016). “Enhancing homology-directed genome editing by catalytically active and inactive CRISPR-Cas9 using asymmetric donor DNA.” Nat Biotechnol 34(3): 339–344.

Riglar, D. T., D. L. Richmond, L. Potvin-Trottier, A. A. Verdegaal, A. D. Naydich, S. Bakshi, E. Leoncini, L. G. Lyon, J. Paulsson and P. A. Silver (2019). “Bacterial variability in the mammalian gut captured by a single-cell synthetic oscillator.” Nat Commun 10(1): 4665.

Santos-Moreno, J., E. Tasiudi, J. Stelling and Y. Schaerli (2020). “Multistable and dynamic CRISPRi-based synthetic circuits.” bioRxiv: 756338.

Scott, S. R., M. O. Din, P. Bittihn, L. Xiong, L. S. Tsimring and J. Hasty (2017). “A stabilized microbial ecosystem of self-limiting bacteria using synthetic quorum-regulated lysis.” Nat Microbiol 2: 17083.

Sheth, R. U., S. S. Yim, F. L. Wu and H. H. Wang (2017). “Multiplex recording of cellular events over time on CRISPR biological tape.” Science 358(6369): 1457–1461.

Shibata, M., H. Nishimasu, N. Kodera, S. Hirano, T. Ando, T. Uchihashi and O. Nureki (2017). “Real-space and real-time dynamics of CRISPR-Cas9 visualized by high-speed atomic force microscopy.” Nat Commun 8(1): 1430.

Singh, D., J. Mallon, A. Poddar, Y. Wang, R. Tippana, O. Yang, S. Bailey and T. Ha (2018). “Real-time observation of DNA target interrogation and product release by the RNA-guided endonuclease CRISPR Cpf1 (Cas12a).” Proc Natl Acad Sci U S A 115(21): 5444–5449.

Singh, D., S. H. Sternberg, J. Fei, J. A. Doudna and T. Ha (2016). “Real-time observation of DNA recognition and rejection by the RNA-guided endonuclease Cas9.” Nat Commun 7: 12778.

Srinivas, N., J. Parkin, G. Seelig, E. Winfree and D. Soloveichik (2017). “Enzyme-free nucleic acid dynamical systems.” Science 358 (6369).

St-Pierre, F., L. Cui, D. G. Priest, D. Endy, I. B. Dodd and K. E. Shearwin (2013). “One-step cloning and chromosomal integration of DNA.” ACS Synth Biol 2(9): 537–541.

Stricker, J., S. Cookson, M. R. Bennett, W. H. Mather, L. S. Tsimring and J. Hasty (2008). “A fast, robust and tunable synthetic gene oscillator.” Nature 456(7221): 516–519.

Toth, E., E. Varga, P. I. Kulcsar, V. Kocsis-Jutka, S. L. Krausz, A. Nyeste, Z. Welker, K. Huszar, Z. Ligeti, A. Talas and E. Welker (2020). “Improved LbCas12a variants with altered PAM specificities further broaden the genome targeting range of Cas12a nucleases.” Nucleic Acids Res 48(7): 3722–3733.

Wang, P., L. Robert, J. Pelletier, W. L. Dang, F. Taddei, A. Wright and S. Jun (2010). “Robust growth of Escherichia coli.” Curr Biol 20(12): 1099–1103.

Yeung, E., A. J. Dy, K. B. Martin, A. H. Ng, D. Del Vecchio, J. L. Beck, J. J. Collins and R. M. Murray (2017). “Biophysical Constraints Arising from Compositional Context in Synthetic Gene Networks.” Cell Syst 5(1): 11–24 e12.

Zetsche, B., J. S. Gootenberg, O. O. Abudayyeh, I. M. Slaymaker, K. S. Makarova, P. Essletzbichler, S. E. Volz, J. Joung, J. van der Oost, A. Regev, E. V. Koonin and F. Zhang (2015). “Cpf1 is a single RNA-guided endonuclease of a class 2 CRISPR-Cas system.” Cell 163(3): 759–771.

Zhang, S. and C. A. Voigt (2018). “Engineered dCas9 with reduced toxicity in bacteria: implications for genetic circuit design.” Nucleic Acids Res 46(20): 11115–11125.

Zhang, Z. B., Q. Y. Wang, Y. X. Ke, S. Y. Liu, J. Q. Ju, W. A. Lim, C. Tang and P. Wei (2017). “Design of Tunable Oscillatory Dynamics in a Synthetic NF-kappaB Signaling Circuit.” Cell Syst 5(5): 460–470 e465.

