## Supporting Information for "Towards a translationally-independent RNA-based synthetic oscillator using deactivated CRISPR-Cas"

#### Methods and Materials

***Plasmid and Strain Construction.*** CRISPR DNA fragments were PCR amplified from existing plasmids: dCas9 system from pdCas9-bacteria, pAN-AND and pAN-OR (Addgene # 62308, 62307, 44249) [1,2]. For dCas12a, Cas12a pieces were PCR amplified with added D917A mutation from pY001 (pFnCpf1_full; Addgene #69973) [3]. Other DNA pieces not obtained by PCR were synthesized as IDT gBlock gene fragments (Integrated DNA Technologies). Plasmids were made by Gibson assembly [4] or restriction enzyme digestion and ligation with T4 ligase in *E. coli* DH5a. Either a high (ColE1), medium (p15A) or low (SC101) copy number origin of replication was used. Genomic integration into the attB site of *E. coli* was performed with the pOSIP plasmid system [5], with clonetegration plasmid cloning done in TransforMax EC100D pir+ electrocompetent *E. coli* (Lucigen).

All mother machine strains were in *E. coli* MG1655 with ΔmotA, a motility knockout for mother machine growth and an integrated constitutive red fluorescent protein reporter for use as a segmentation marker (strain NDL162) [6].

***Mother Machine and Microscopy.*** Mother machine microfluidic chips made of polydimethylsiloxane (PDMS) were cast from a patterned silicon wafer and bonded to a No. 1.5 glass coverslip (Fisher Scientific), as previously described [6,7]. Most runs used chips with trench dimensions of 1.5 µm wide and high, with different lengths (10+ µm) used throughout. EZ Rich Defined Medium (EZ-RDM, TekNova) was used with 0.85 g/L Pluronic F108 (Sigma-Aldrich) to prevent cell adhesion to PDMS. Runs did not have supplemented antibiotic. For induction, typical concentrations were 1.88 ng/mL anhydrotetracycline (aTc) and 50 µM 2,4-diacetylphloroglucinol (DAPG). For most runs, media was flown at 20 µL/min through Tygon microfluidic tubing (US Plastic Corp 56515) with NE-300 syringe pumps (New Era Pump Systems).

Cells were imaged on modified Nikon Ti2-E inverted microscopes with Plan Apo 40x air objective (NA 0.95, Nikon), Perfect Focus System (PFS, Nikon), Nikon Ei-S-ER motorized stage with encoders, and high-resolution CMOS camera (Andor Zyla 4.2 PLUS), all controlled with NIS-Elements software (Nikon). All runs were kept at 37°C with a temperature-controlled cage incubator (Okolab). Exposure times varied per experimental run, but RFP was imaged at

100-200 ms and YFP for 10-50 ms using an LED excitation system (SPECTRA X light engine, Lumencor). Images were acquired at frame rates of 6-10 min, as noted throughout.

***Image processing and analysis.*** Microscopy images were processed using ImageJ and MATLAB to segment mother cells using code originally developed by Somenath Bakshi. Further analysis was performed in MATLAB with code from S. Bakshi, J. Kuo and R. Yuan. Autocorrelation function (ACF) and power spectrum of the windowed ACF were calculated in MATLAB and as described previously [7]. Interpeak distances and peak heights were determined using MATLAB *findpeaks* function to find local maxima on a Savitzky-Golay filter smoothed signal. After minimal smoothing by fitting a polynomial order 5 with an 11-frame window using MATLAB function *sgolayfilt*, the peak distances were found using MATLAB function *findpeaks*. The minimum peak prominence value was adjusted if the fluctuation heights were not captured based on visual inspection. Inducible cascade measurements plotted as average plus shaded region standard deviation used MATLAB function *confplot* from Michele Giugliano (Brain Mind Institute, EPFL, Lausanne, Switzerland).

***Bulk strain analysis: plate reader and agarose pads.*** Strains were evaluated in 200 µL test cultures in a 96-well flat-bottom black with clear bottom fluorescence plates (Corning 3631) with low evaporation lids (Corning 3931). Plates were grown in a BioTek Synergy H1 plate reader with continuous shaking at 37ºC, including a higher gradient temperature above to reduce condensation. Seed cultures were grown overnight in a separate 96-well plate. Agarose pads to qualitatively look at cultures were prepared with 1% agarose with microscope slides [8].

### Supplementary Figures

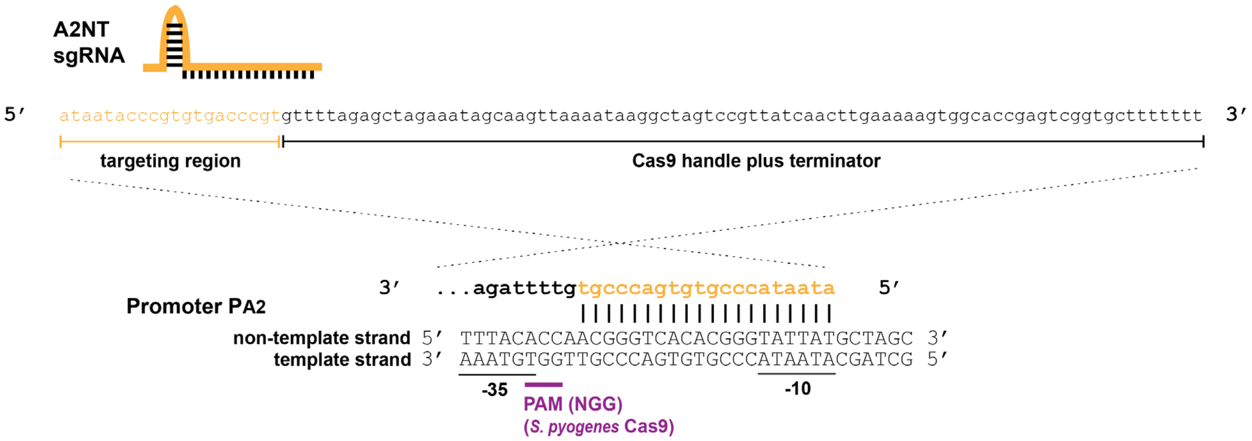

**Figure S1. Single guide RNA (sgRNA) targeting of promoter at the sequence level.** The dCas9-sgRNA complex targets a promoter region between the -35 and -10 boxes. In this case, A2NT sgRNA targets the non-template strand of Promoter PA2 in the inter-region of the -10 and -35 boxes. The *S. pyogenes* dCas9 is targeted to "NGG" protospacer adjacent motif (PAM) site. Other sgRNA and promoter pair designs target the same regions with different sequences for the -35 and -10 inter-region. Parts are from [2].

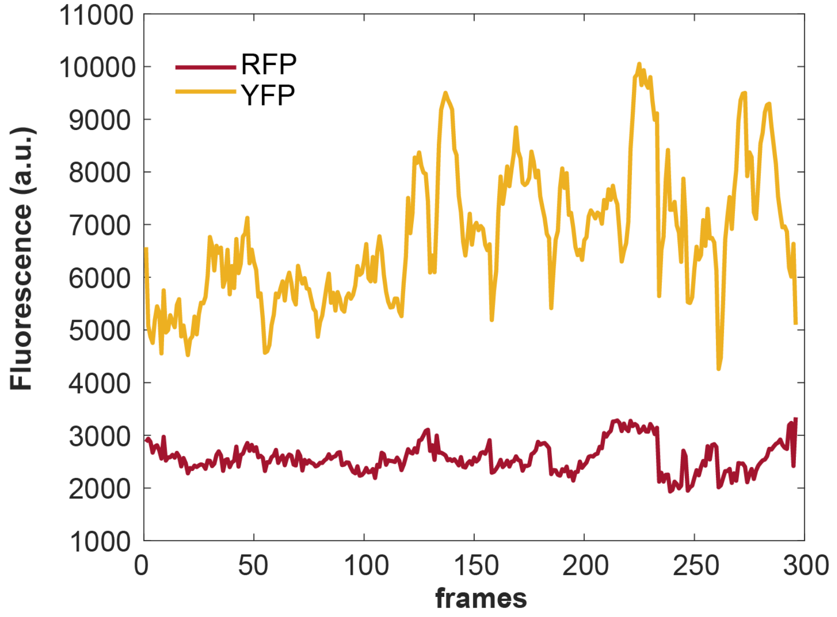

**Figure S2. Mother machine reporter and segmentation traces for dCas9 RNA oscillator, one-plasmid form.** RFP segmentation marker (modified mKate2) and YFP reporter (mVenus) traces from mother machine run for one-plasmid dCas9 oscillator (JK412) cell in mother machine. EZ-RDM, 8 min/frame. Significant fluctuations are from YFP.

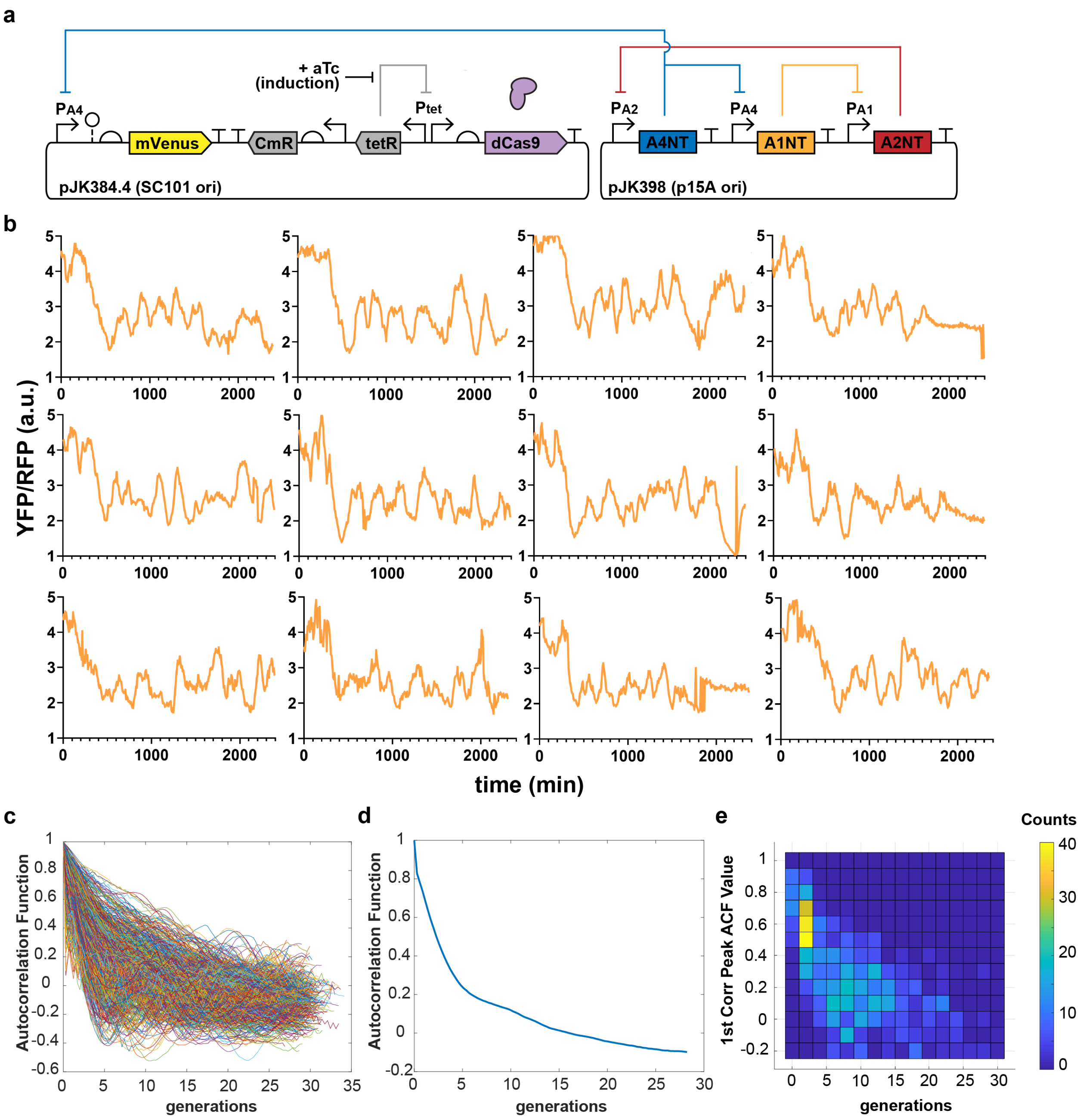

**Figure S3. Two-plasmid dCas9 RNA oscillator with PA4 promoter.** (a) Diagram of circuit on two plasmids, with PA4-mVenus reporter. (b) Example mother cell traces from mother machine run in EZ-RDM with 0.625 ng/mL aTc added at ~280 min and changed to 1.3 ng/mL aTc at ~2000 min. Mean generation time ~28 min. Acquired every 8 minutes. (c) Autocorrelation function (ACF) of 887 individual cells. (d) Population-averaged ACF. (e) Individual first correlation peaks have a wide distribution.

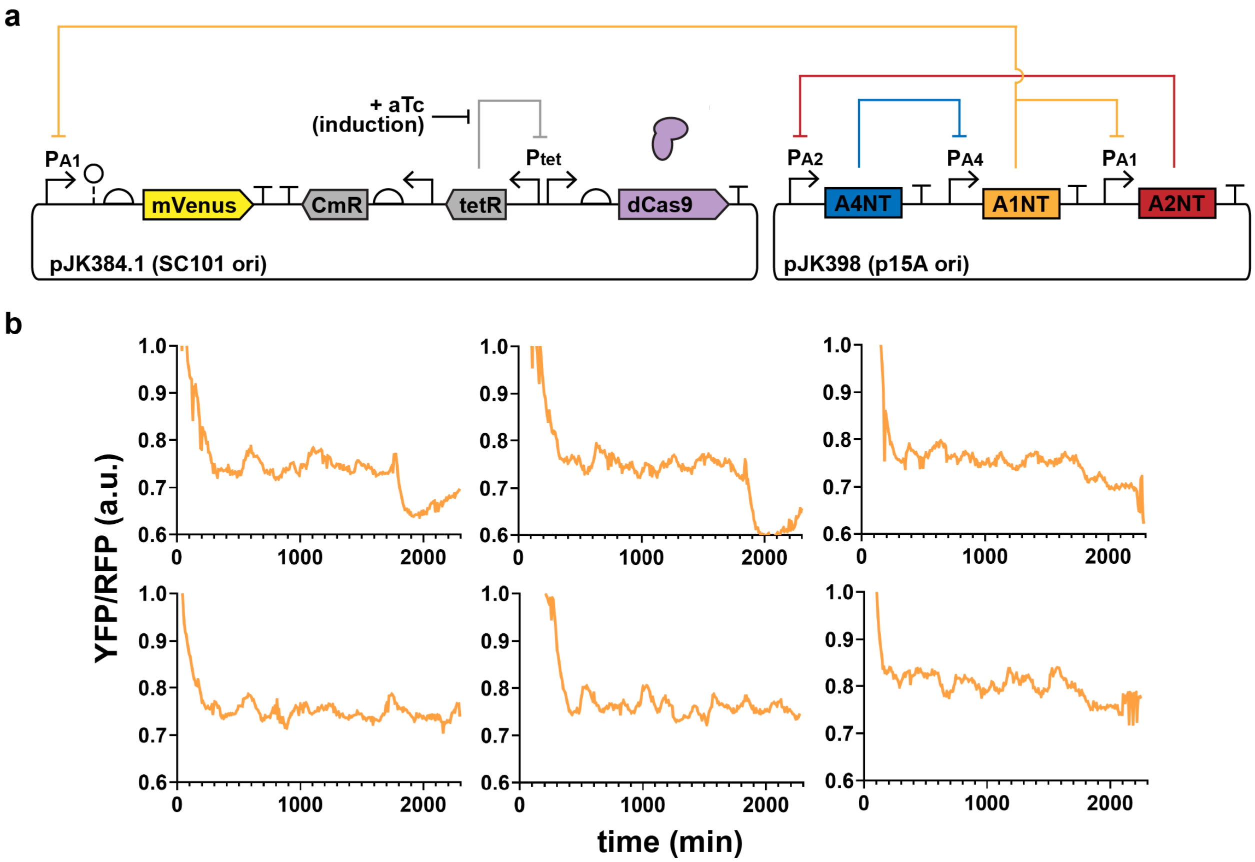

**Figure S4. Two-plasmid dCas9 RNA oscillator with PA1 reporter.** (a) Diagram of circuit on two plasmids, with PA1-mVenus reporter. (b) Example mother cell traces from mother machine run in EZ-RDM with 0.625 ng/mL aTc added ~142 min. Median generation time ~26.5 min. Acquired every 8 minutes.

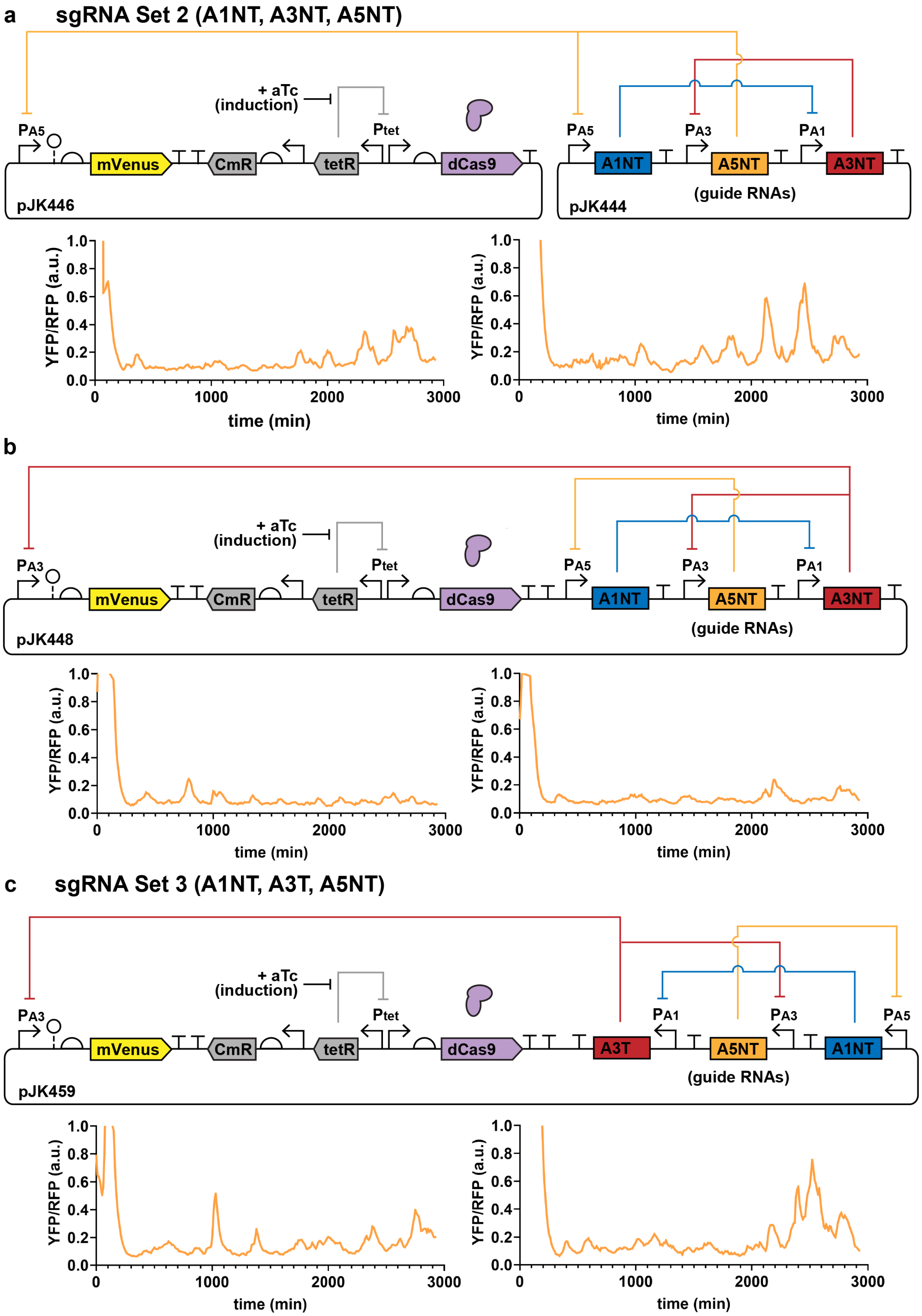

**Figure S5. Other guide RNA sets for dCas9 oscillator.** Designs with different sets of orthogonal sgRNAs [2]. Signals were repressed to low levels after aTc induction at 90 min. Some fluctuations were preserved, as shown in the example cell traces, but signal-to-noise was too low for many peaks to characterize. Cells grown in mother machine chip with EZ-RDM. Average generation times ~27 min. Acquired every 10 min.

##### *Attempted Synchronization*

We tried to synchronize dCas9 RNA oscillator fluctuations across a population by adding an accessory plasmid with an inducible extra guide RNA (Fig. S6). Previous systems likely had high levels of each sgRNA to start. The intended effect was for all cells to have an initial high amount of the induced sgRNA relative to other sgRNAs, to strongly bias cells into that starting state. This could also reduce cell fluctuations between guide RNA states initially because of similar levels of each sgRNA. Neither induction overnight or for a few hours before dCas9 induction changed performance when compared to the genome integrated dCas9 oscillator circuit (JK422) without an extra sgRNA plasmid, suggesting little effect of the extra initial sgRNA.

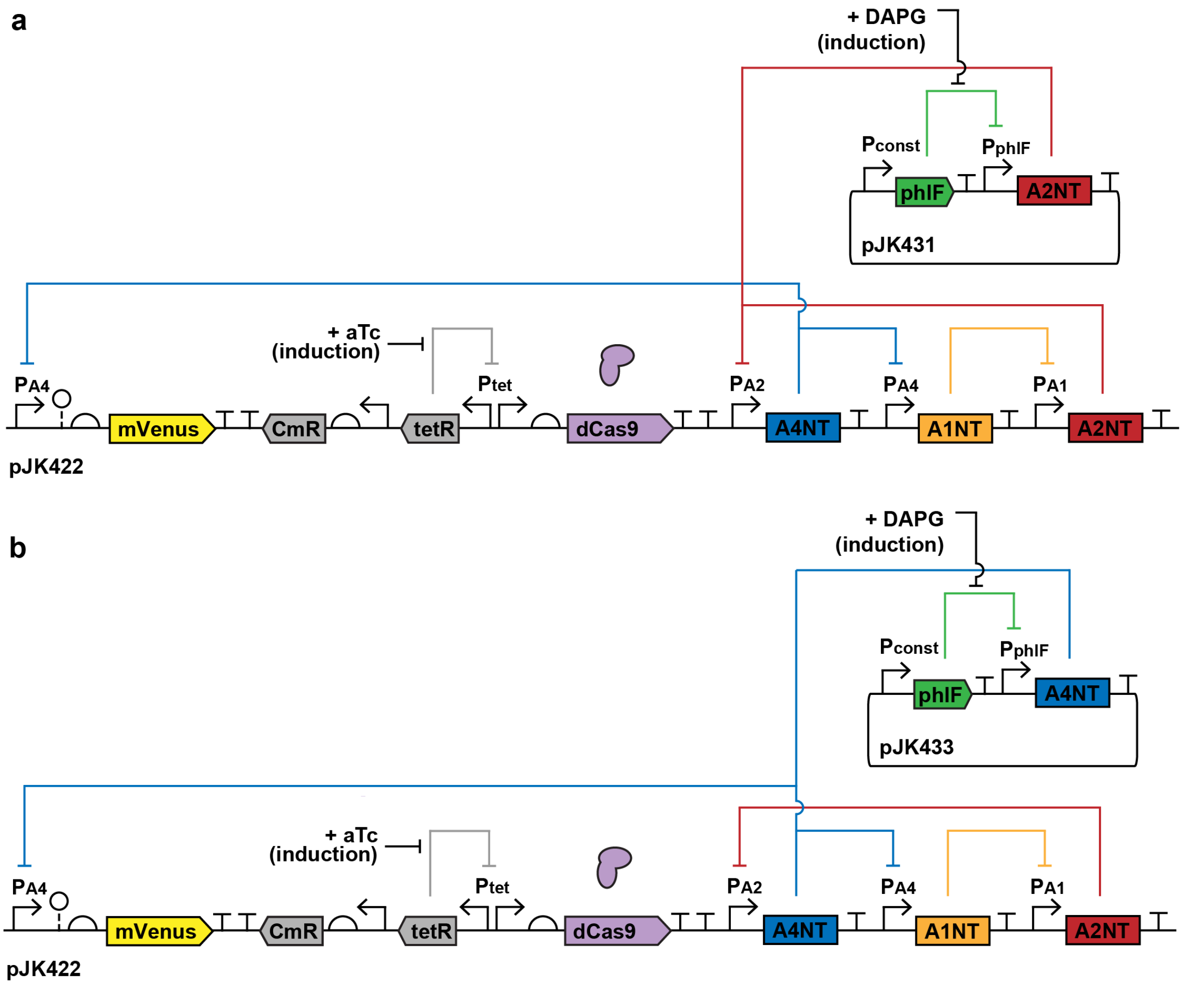

**Figure S6. Genomic dCas9 RNA oscillator (JK422) plus accessory sgRNA plasmid.** JK422 strains had an accessory plasmid with small molecule DAPG-inducible production of one of the sgRNAs. The intention was to help maintain cells in one RNA state prior to dCas9 induction, for population synchronization. pJK433 plasmid was also transformed into JK423.

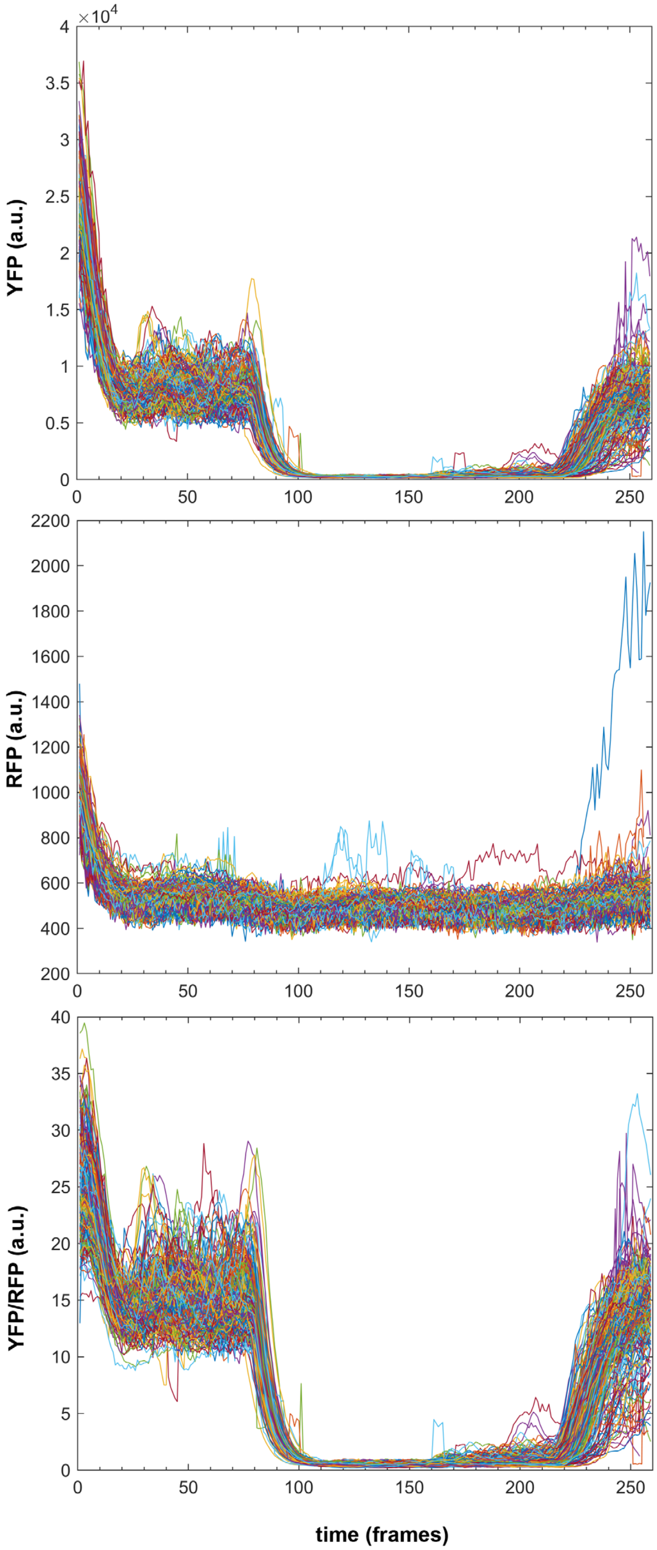

**Figure S7. Signals for YFP and RFP.** Response times for one-plasmid dCas9 cascade (pJK430) shown in Fig. 3b. In the pre-DAPG sgRNA induction phase (frames 20-70), the YFP reporter signals span roughly 0.5 -1x10$^4$, which leads to the 10-20 YFP/RFP signals. Traces shown for 388 cells. 6 min/frame.

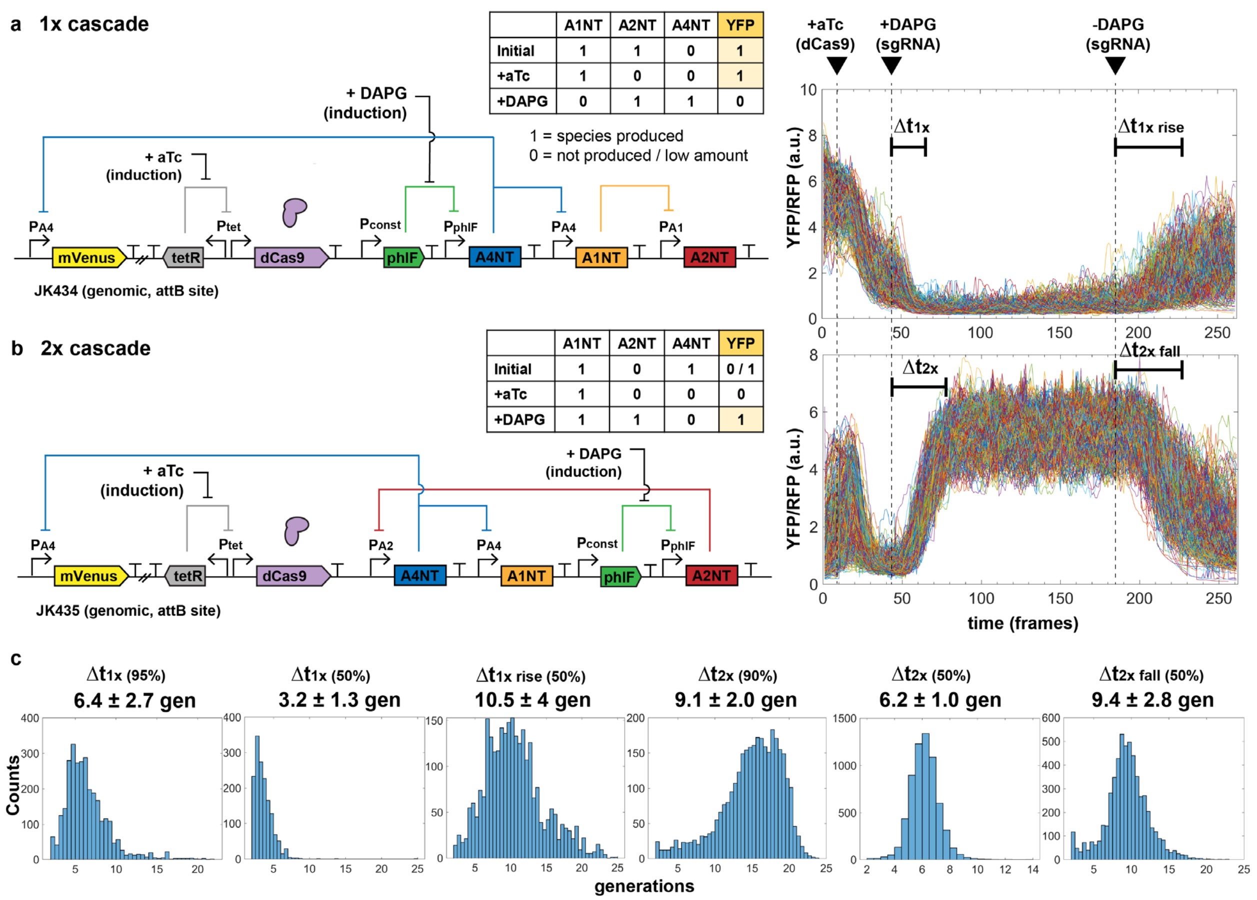

**Figure S8. dCas9 cascade response plots for genomic versions.** Analogous plot as Fig. 3 for genomic versions of dCas9 cascade step circuits. Times are similar, with lower signals as expected. 6 min/frame. Distributions were measured from (a) 2988 cells for pJK434 and (b) 6379 cells for pJK435.

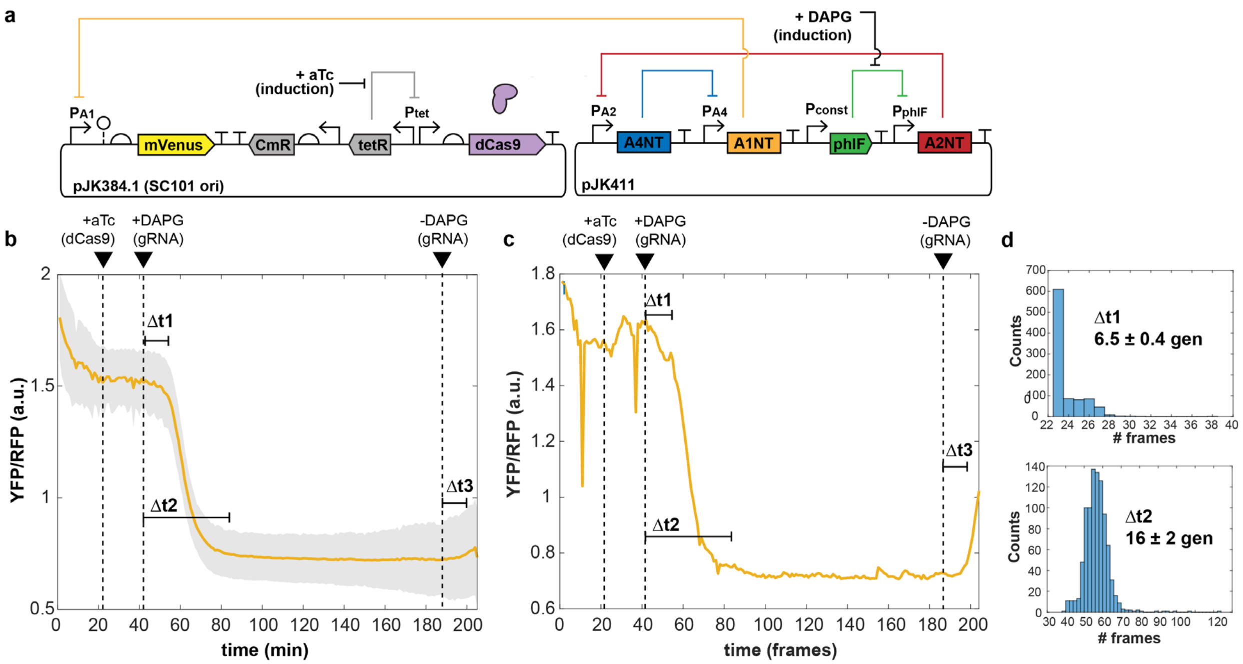

**Figure S9. Response times for two-plasmid dCas9 sgRNAs.** (a) Two-plasmid 3x cascade diagram. Upon DAPG-induced production of A2NT sgRNA, two more steps are needed for A1NT sgRNA repression of mVenus YFP signal. (b) Average reporter fluorescence trace of 1048 cells, with shaded region showing standard deviation. (c) Single trace of representative mother cell, highlighting fluorescence recovery at end upon DAPG removal. (c) Histograms of characteristic time lengths for each step.

***dCas9 Degradation***

Since dCas9 binding was previously shown to be very tight, with off rates longer than *E. coli* division times [9], we thought dCas9 protein degradation might be important. However, tests of adding four ssrA-based (small stable RNA A) tags [10] C-terminal to dCas9 did not noticeably improve behavior. These ssrA tags have the sequence “AADENYALAA”, where the underlined portion varies to determine strength. The strong tags (“LVA” and “LAA”) reduced dCas9 repression in a plate reader bulk assay, as did a medium strength tag (“AAV”), while the weakest tag (“ASV”) performed similarly to without degradation (Fig. S10). Mother machine runs also did not show improved performance, with the strongest LAA tag abolishing fluctuations. Because of this, we used an untagged dCas9 in our other designs.

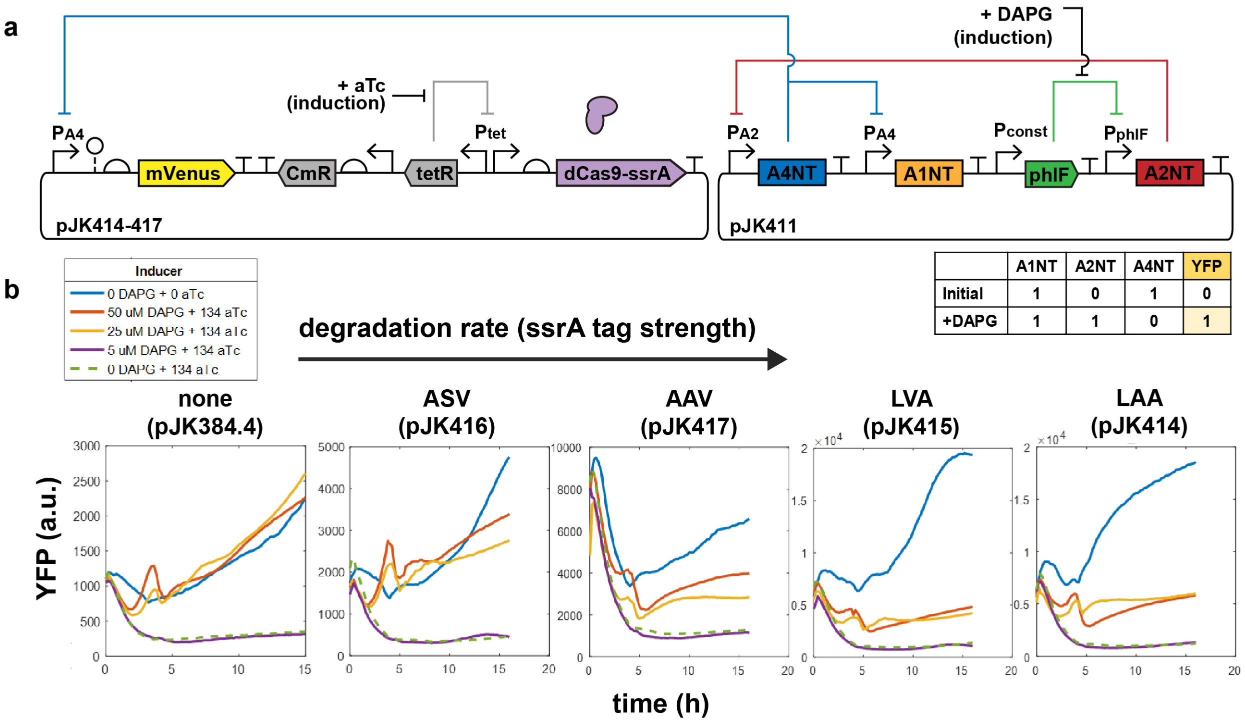

**Figure S10. Degradation tag effects on dCas9 in bulk plate reader assay.** (a) An inducible 3x sgRNA cascade was used with a PA4-mVenus reporter to test dCas9 constructs with C-terminal ssrA degradation tags (ex., dCas9 + AADENYALAA). (b) Responses of YFP signal in system with added degradation tags to dCas9. Tags are shown in order of increasing putative strength [10], from none to ASV, AAV, LVA and LAA. Inducer added was 0-50 µM DAPG and 134 nM (62.5 ng/mL) aTc. No induction (blue line) allowed free mVenus reporter production, while dCas9 induction without sufficient sgRNA production (purple and dashed green lines) repressed mVenus. DAPG-induced sgRNA production led to repression of promoter PA2 and A4NT sgRNA, allowing mVenus expression (red and orange lines) to varying degrees depending on dCas9 degradation tag.

##### *aTc Titration*

In addition to adjusting aTc levels during mother machine runs, plate reader assays helped determine aTc effects. Looking at different YFP reporter promoters with different induced sgRNAs showed even low amounts of aTc could be effective for repression (Fig. S11).

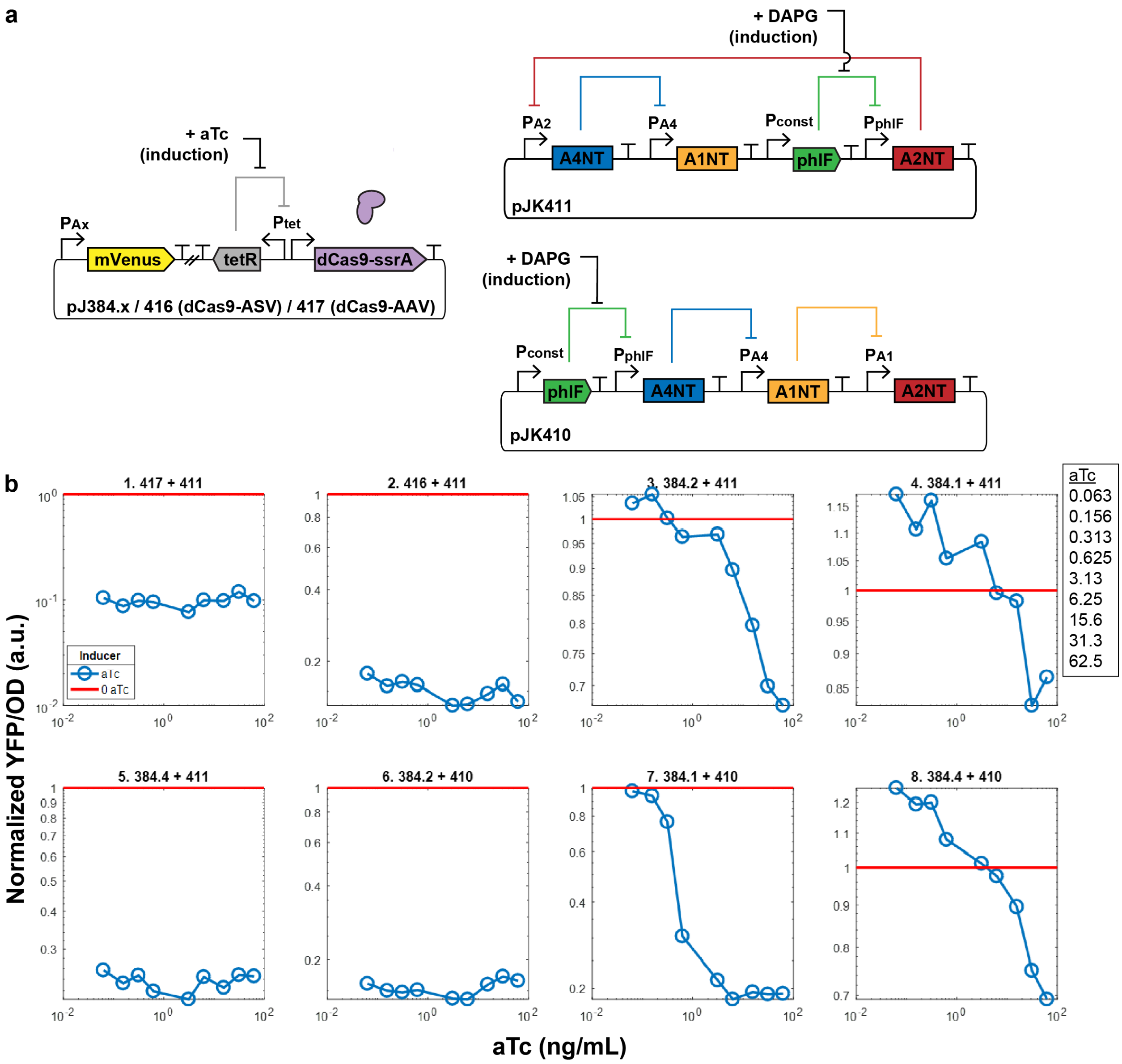

**Figure S11. Inducer aTc titration for dCas9 expression in bulk plate reader assay.** (a) Inducible sgRNA cascades were used with a varying PAx-Venus reporters (384.1,2,4 for PA1, PA2, PA4) to produce constant levels of sgRNAs. pJK416 and pJK417 both have PA4 promoter for mVenus, with added ssrA tags ASV and AAV on dCas9. (b) Plate assays were run. Red line shows level without aTc added (aTc = 0). All values are normalized to aTc = 0 for that strain. Concentrations of aTc, increasing order (ng/uL): 0.063, 0.156, 0.313, 0.625, 3.13, 6.25, 15.6, 31.3, 62.5. DAPG to induce sgRNA was not added. Without DAPG added, pJK410+411 act as constitutive producers of sgRNAs.

In Fig. S11b panels #1,2,5,6, even the lowest level of aTc (0.063 ng/mL) was enough dCas9 production to provide repression comparable to 1000x more aTc (62.5 ng/mL). In panel #7, a clearer response curve was seen for PA1 repression, with similar repression from 62.5 down to 3.13 ng/mL but a drop in effectiveness at lower concentrations.

Along with these results, we noticed higher toxicity to cells in mother machine runs at higher aTc amounts. Because of this, we tried to use a low amount but enough such that degradation over the 24+ of growth and imaging would not be a significant factor. A balance of this was 0.625 ng/mL.

*Testing dCas9 induction levels in the mother machine.* After previous bulk plate reader assays showed a lower amount of dCas9 may be helpful, we used these lower aTc levels to test in single-cell mother machine experiments. When using 62.5 ng/mL aTc for induction, cells showed signs of stress – much higher filamentation rates and halting of growth. Lowering aTc 100-fold to 0.625 ng/mL, fluctuations persisted, but somewhat better performance was observed at three-fold higher to 1.88 ng/mL (Fig. S12). When further raised to 6.25 ng/mL aTc, cells again showed the greater toxicity effects seen at 62.5 ng/mL, with an increased rate of halted growth while maintaining high levels of YFP reporter.

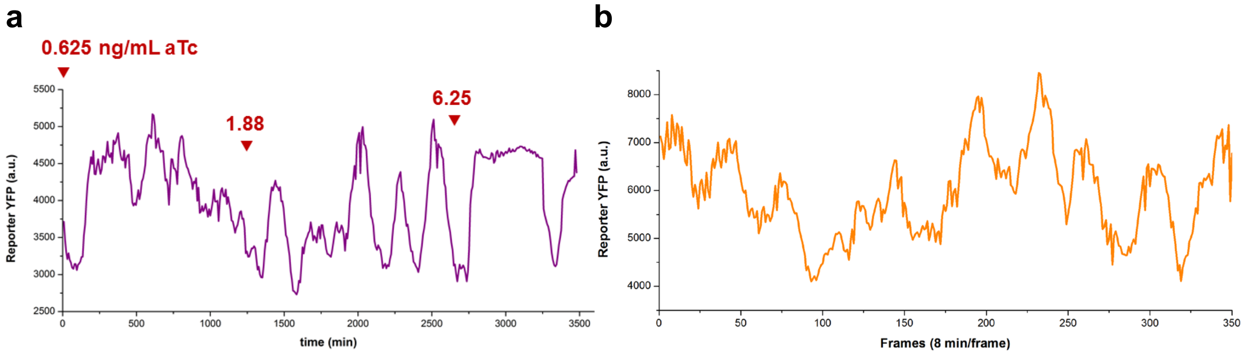

**Figure S12. aTc titration in mother machine.** Following a cross-section of a mother cell, the reporter YFP signal fluctuates over time. One-plasmid dCas9 oscillator (JK412) cells grown in mother machine chips with EZ-RDM. (a) Marked timepoints are where aTc concentration was changed to the indicated amount (ng/mL). Prior to addition of aTc for dCas9 induction, cells had a consistently high level of fluorescence. (b) In a separate run, dCas9 induction was held constant at 1.88 ng/mL. A 350-frame (~46 h) window of a 96+ h run is shown of a representative trace of a clearly fluctuating cell.

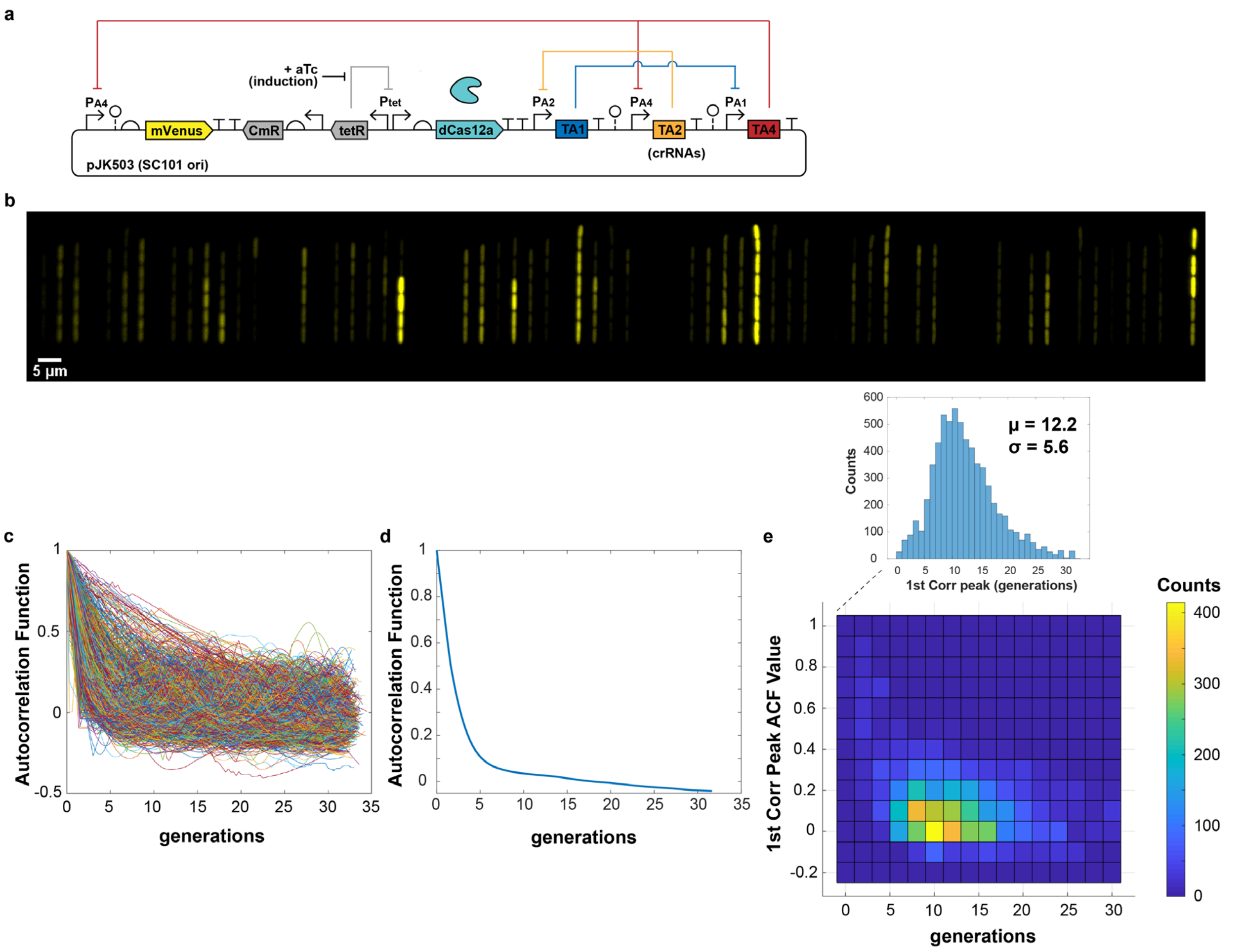

**Figure S13. One-plasmid dCas12a RNA oscillator, with crRNAs in same transcriptional direction as dCas12a (JK503).** (a) Diagram of 1-plasmid dCas12a RNA oscillator design with dCas12a and crRNAs in same transcriptional direction (pJK503). (b) Snapshot of *E. coli* MG1655 cells with plasmid in mother machine. Cells show variation, with strong repression. (c) Individual autocorrelation function (ACF) traces. (d) The population-averaged ACF shows a slower decay. (e) Individual first correlation peaks have a wide distribution. Measured from 6653 cells.

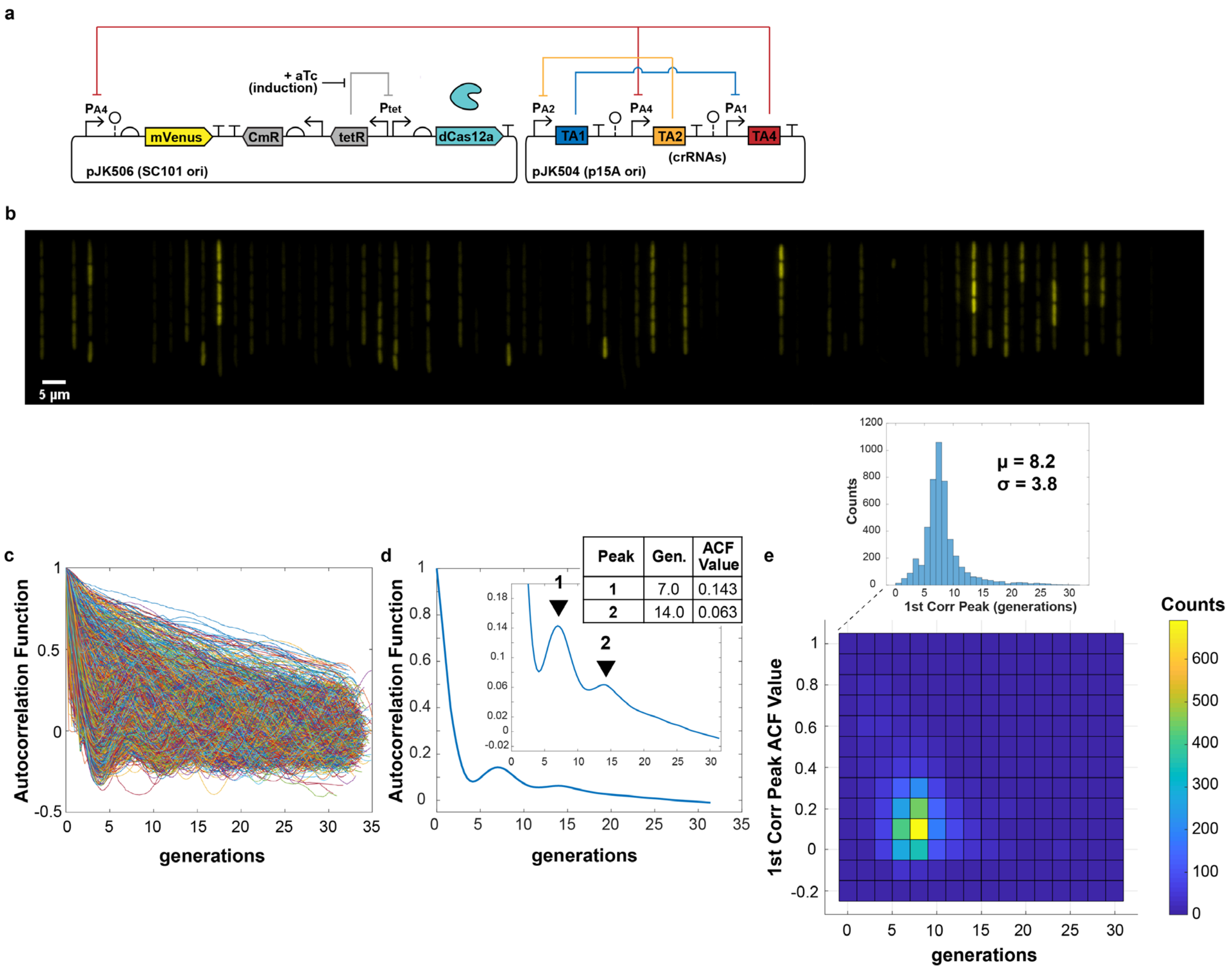

**Figure S14. Two-plasmid dCas12a RNA oscillator (JK504+506).** (a) Diagram of 2-plasmid dCas12a RNA oscillator design (pJK504 + pJK506). (b) Snapshot of *E. coli* MG1655 cells with plasmids in mother machine. Cells show variation, with strong repression. (c) Individual autocorrelation function (ACF) traces. (d) The population-averaged ACF showed distinct correlation peaks. (e) Individual first correlation peaks had a narrower distribution. Measured from 4962 cells.

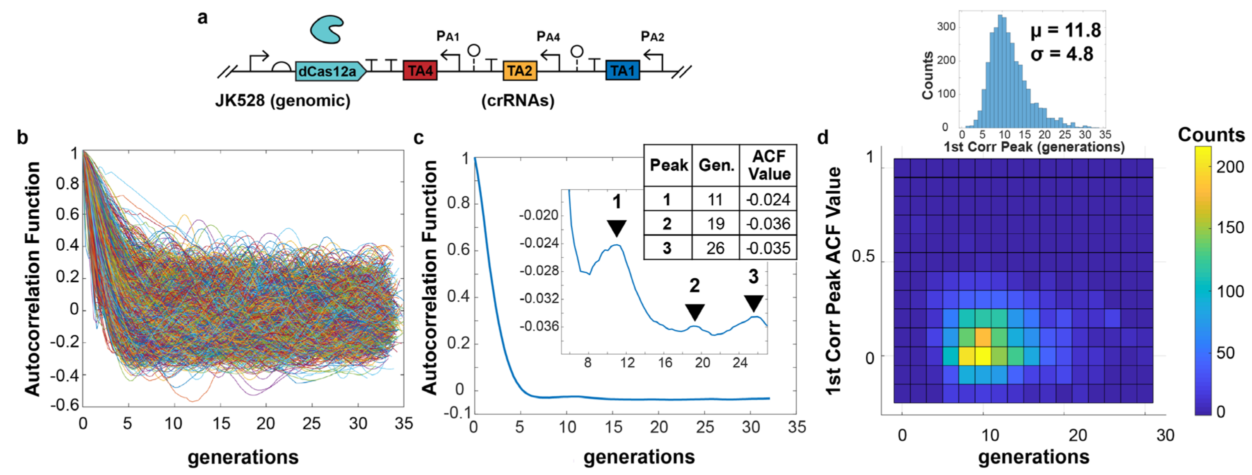

**Figure S15. Genome-integrated dCas12a RNA oscillator (JK528).** (a) Diagram of genome-integrated dCas12a RNA oscillator design (JK528), with transcriptionally convergent crRNAs. (b) Individual autocorrelation function (ACF) traces. (c) The population-averaged ACF showed distinct correlation peaks, though small. (d) Individual first correlation peaks had a wide distribution. Measured from 6636 cells.

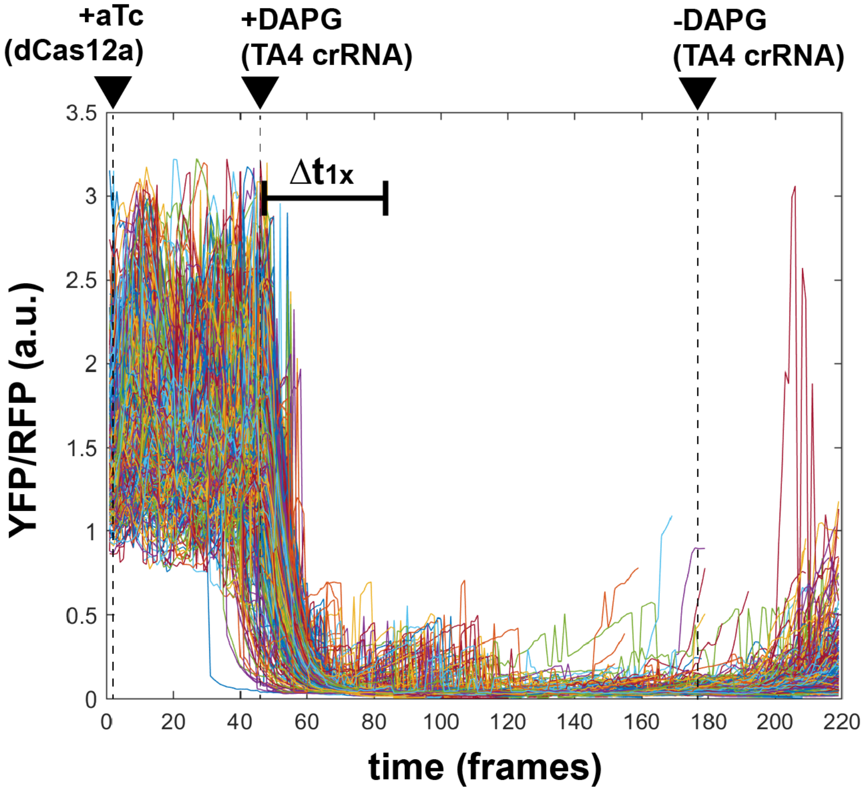

**Figure S16. dCas12a crRNA response times single cells.** Individual cell traces for dCas12a 1x (pJK526). 1049 cells. Average shown in main text Fig. 6b.

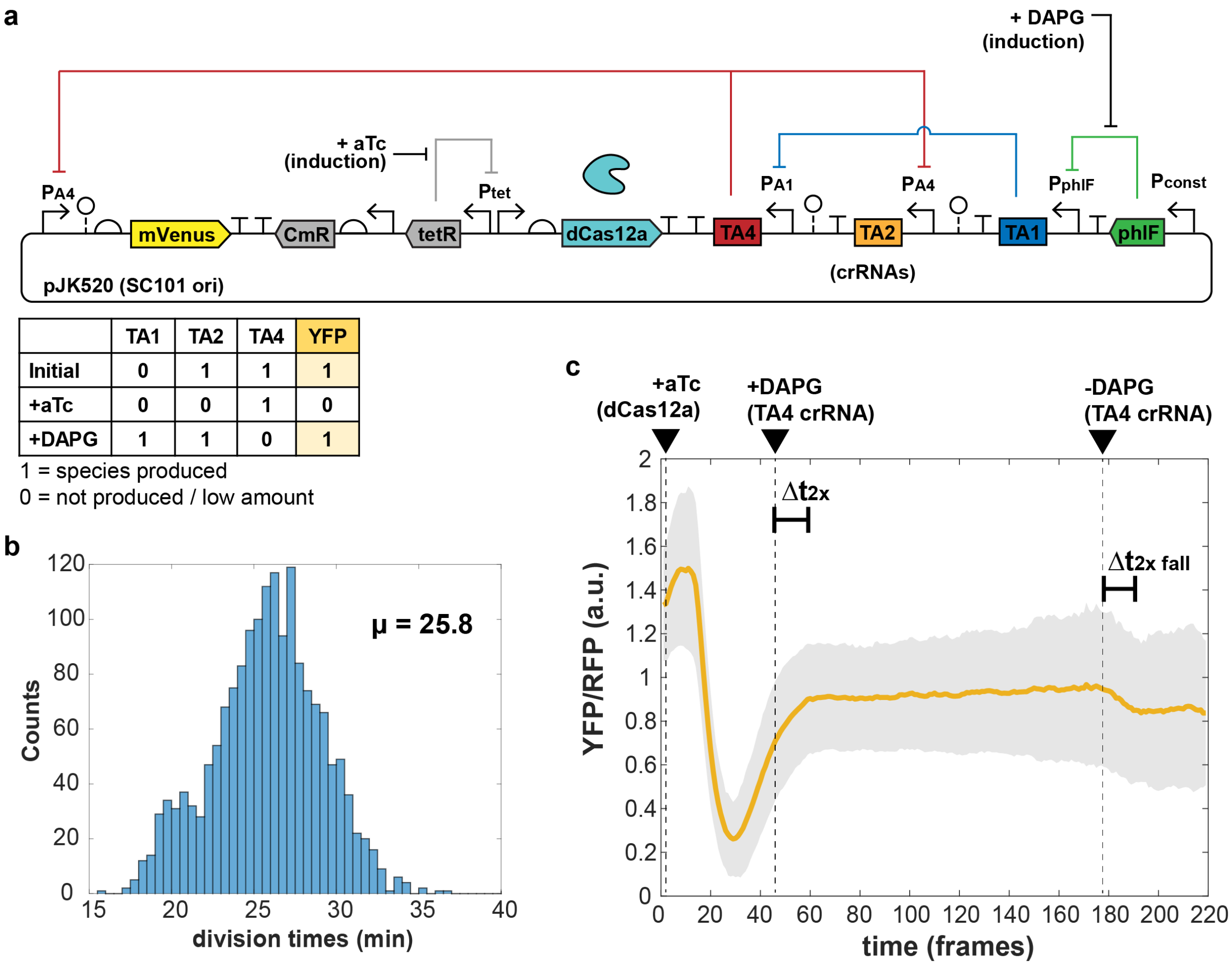

| | TA1 | TA2 | TA4 | YFP |
|---|---|---|---|---|
| Initial | 0 | 1 | 1 | 1 |
| +aTc | 0 | 0 | 1 | 0 |
| +DAPG | 1 | 1 | 0 | 1 |

**Figure S17. dCas12a crRNA dCas12a crRNA two-step response times**. (a) A single constitutive promoter of the free-running dCas12a oscillator was replaced with a PhlF repressor-based promoter system, allowing inducible control with small molecule DAPG. *E. coli* MG1655 cells with plasmid were grown in EZ-RDM in the mother machine. (b) Histogram of average division times. (c) Averaged trace from 1698 cells of plasmid version (pJK520), with shaded region showing one standard deviation of the averaged values. 1 frame = 6 minutes.

#### Supplementary Videos

**Video S1. dCas9 RNA oscillator (JK412) in mother machine.**
*E. coli* MG1655 cells with dCas9 RNA oscillator plasmid (pJK412) are in a mother machine chip with EZ-RDM for exponential growth. YFP reporter fluorescence shown, 8 min per frame. Inducer aTc begins at 0.625 ng/mL and is changed to 1.88 ng/mL at time 20:00.

**Video S2. dCas12a RNA oscillator (JK511) in mother machine.**
*E. coli* MG1655 cells with dCas12a RNA oscillator plasmid (pJK511) are in a mother machine chip with EZ-RDM for exponential growth. YFP reporter fluorescence shown, 8 min per frame. Inducer aTc (1.88 ng/mL) added at video start.

**Video S3. dCas12a RNA oscillator (JK503) in mother machine.**
*E. coli* MG1655 cells with dCas12a RNA oscillator plasmid (pJK503, which has crRNAs in the same transcriptional direction as dCas12a) are in a mother machine chip with EZ-RDM for exponential growth. YFP reporter fluorescence shown, 8 min per frame. Inducer aTc (1.88 ng/mL) added at time 2:56.

**Video S4. dCas12a RNA oscillator as two plasmids (pJK504+506) in mother machine.**
*E. coli* MG1655 cells with dCas12a RNA oscillator plasmids (pJK504+506) are in a mother machine device with EZ-RDM for exponential growth. YFP reporter fluorescence shown, 8 min per frame. Inducer aTc (1.88 ng/mL) added at time 2:56.

### Supplementary Tables

**Table S1. Plasmids for dCas9 designs.** ORI: origin of replication. Res: antibiotic resistance: Cm, chloramphenicol; Amp, ampicillin; K, kanamycin.

| Name | Description | ORI | Res |
|---|---|---|---|
| pJK384.4 | dCas9 + PA4-mVenus | SC101 | Cm |
| pJK384.1 | dCas9 + PA1-mVenus | SC101 | Cm |
| pJK384.2 | dCas9 + PA2-mVenus | SC101 | Cm |
| pJK388 | full dCas9 sgRNA oscillator v1 (dCRv1) (ColE1) | ColE1 | Amp |
| pJK398 | full dCas9 sgRNA oscillator v1 (dCRv1) (p15A) | p15A | Amp |
| pJK410 | PphlF_A4NT in dCRv1 | p15A | Amp |
| pJK411 | PphlF_A2NT in dCRv1 | p15A | Amp |
| pJK412 | dCas9 oscillator v1 (dCRv1) + [dCas9 + PA4-mVenus] | SC101 | Cm |
| pJK413 | dCas9 oscillator v1 (dCRv1) + [dCas9 + PA4-mVenus] (sgRNAs flip of 412) | SC101 | Cm |
| pJK414 | dCas9-LAA + PA4-mVenus | SC101 | Cm |
| pJK415 | dCas9-LVA + PA4-mVenus | SC101 | Cm |
| pJK416 | dCas9-ASV + PA4-mVenus | SC101 | Cm |
| pJK417 | dCas9-AAV + PA4-mVenus | SC101 | Cm |
| pJK418 | dCas9-LAA + PA1-mVenus | SC101 | Cm |
| pJK419 | dCas9-LVA + PA1-mVenus | SC101 | Cm |
| pJK421 | dCas9-AAV + PA1-mVenus | SC101 | Cm |
| pJK422 | dCas9 oscillator v1 (dCRv1) + [dCas9 + PA4-mVenus]. (Integration of 412; pOSIP) | R6K gamma | Cm/K |
| pJK423 | dCas9 oscillator v1 (dCRv1) + [dCas9 + PA4-mVenus]. (Integration of 413; pOSIP) | R6K gamma | Cm/K |
| pJK424 | [PA4-mVenus]. (Integration; pOSIP) | R6K gamma | Cm/K |
| pJK425 | dCRv1 sgRNAs only (412 direction). (Integration; pOSIP) | R6K gamma | K |
| pJK426 | dCRv1 sgRNAs only (413 direction). (Integration; pOSIP) | R6K gamma | K |
| pJK427 | [PA4-mVenus + dCas9]. (Integration; pOSIP) | R6K gamma | Cm/K |
| pJK428 | TetR-dCas9. (Integration; pOSIP) | R6K gamma | K |
| pJK429 | PphlF_A4NT in dCRv1, 1x cascade + [dCas9 + PA4-mVenus] | SC101 | Cm |
| pJK430 | PphlF_A2NT in dCRv1, 2x cascade + [dCas9 + PA4-mVenus] | SC101 | Cm |
| pJK431 | PphlF-A2NT (p15A ori) | p15A | Amp |
| pJK433 | PphlF-A4NT (p15A ori) | p15A | Amp |
| pJK434 | PphlF_A4NT, 1x cascade + [dCas9 + PA4-mVenus]. (Integration of 429; pOSIP) | R6K gamma | Cm/K |
| pJK435 | PphlF_A2NT, 2x cascade + [dCas9 + PA4-mVenus]. (Integration of 430; pOSIP) | R6K gamma | Cm/K |
| pJK436 | PphlF_A1NT in dCRv1 | p15A | Amp |
| pJK437 | PphlF_A1NT in dCRv1 + [dCas9 + PA4-mVenus] | SC101 | Cm |
| pJK439 | PphlF_A1NT in dCRv1 + [dCas9 + PA4-mVenus]. (Integration of 437; pOSIP) | R6K gamma | Cm/K |
| pJK444 | dCRv2 (PA5-A1NT + PA3-A5NT + PA1-A3NT) in p15A | p15A | Amp |
| pJK446 | dCas9 + PA3-mVenus | SC101 | Cm |
| pJK447 | dCRv3 (PA5-A1NT + PA3-A5NT + PA1-A3T) in p15A | p15A | Amp |
| pJK448 | dCRv2 + [dCas9 + PA3-mVenus] | SC101 | Cm |
| pJK458 | dCRv2 flip + [dCas9 + PA3-mVenus] | SC101 | Cm |
| pJK459 | dCRv3 flip + [dCas9 + PA3-mVenus] | SC101 | Cm |

**Table S2. Plasmids for dCas12a designs.** ORI: origin of replication. Res: antibiotic resistance: Cm, chloramphenicol; Amp, ampicillin; K, kanamycin.

| Name | Description | ORI | Res |
|---|---|---|---|
| pJK503 | dCas12a oscillator, same direction crRNAs. dCas12a (D917A) + PA4-mVenus | SC101 | Cm |
| pJK504 | dCas12a crRNAs | p15A | Amp |
| pJK506 | dCas12a (D917A) + PA4-mVenus. (removed crRNAs) | SC101 | Cm |
| pJK507 | dCas12a crRNAs. (flipped middle crRNA) | p15A | Amp |
| pJK511 | dCas12a oscillator, reverse direction crRNAs. dCas12a (D917A) + PA4-mVenus | SC101 | Cm |
| pJK517 | PphlF-TA4 in crRNAs | p15A | Amp |
| pJK520 | PphlF-TA1 crRNAs, 2x cascade. dCas12a (D917A) + PA4-mVenus. | SC101 | Cm |
| pJK526 | PphlF-TA4 crRNAs, 1x cascade. dCas12a (D917A) + PA4-mVenus. | SC101 | Cm |
| pJK528 | dCas12a oscillator, reverse crRNAs + PA4-mVenus. (Integration of 511; pOSIP) | R6K gamma | Cm/K |

**Table S3. Strains.**

| Name | Description |
|---|---|
| NDL162 | *E. coli* MG1655 ΔmotA PRNA1-mKate2 (ref. [6]) |
| | (mKate2 N-term 4 amino acids replaced by mCherry N-term 11 amino acids) |
| JK422 | NDL162 attB::dCas9 oscillator v1 [dCRv1 + dCas9 + PA4-mVenus] |
| JK423 | NDL162 attB::dCas9 oscillator v1 [dCRv1 flip sgRNAs + dCas9 + PA4-mVenus] |
| JK434 | PphlF_A4NT in dCRv1, 1x cascade + [dCas9 + PA4-mVenus] |
| JK435 | PphlF_A2NT in dCRv1, 2x cascade + [dCas9 + PA4-mVenus] |
| JK528 | NDL162 attB::dCas12a oscillator, reverse crRNAs + PA4-mVenus. |

**Table S4. Sequences.** (For full plasmid maps/sequences, see Addgene.)

| Name | Description | DNA Sequence (5' to 3') |
|---|---|---|
| mVenus | RiboJ ribozyme + RBS + mVenus + rrnB T1 terminator | Agctgtcaccggatgtgctttccggtctgatgagtccgtgaggacgaaacagcctctacaaataattt<br>tgtttaactttacaaagaggagaaaggtaccgcatgagtaaaggagaagaacttttcactggagttgt<br>cccaattcttgttgaattagatggtgatgttaatgggcacaaattttctgtcagtggagagggtgaag<br>gtgatgcaacatacggaaaacttacccttaaattgatttgcactactggaaaactacctgttccatgg<br>ccaacacttgtcactactttgggttatggtgttcaatgctttgcgagatacccagatcatatgaaaca<br>gcatgactttttcaagagtgccatgcccgaaggttatgtacaggaaagaactatatttttcaaagatg<br>acgggaactacaagacacgtgctgaagtcaagtttgaaggtgatacccttgttaatagaatcgagtta<br>aaaggtattgattttaaagaagatggaaacattcttggacacaaattggaatacaactataactcaca<br>caatgtatacatcacggcagacaaacaaaagaatggaatcaaagcgaacttcaaaattagacacaaca<br>ttgaagatggaggtgttcaactagcagaccattatcaacaaaatactccaattggcgatggccctgtc<br>cttttaccagacaaccattacctgtcctaccaatctaagctttcgaaagatcccaacgaaaagagaga<br>ccacatggtccttcttgagtttgtaacagctgctgggattacacatggcatggatgaactatacaaat<br>aagcttgatatcgaattcctgcagcccgggggatcccattgtacgcgtgctagaggcatcaaataaaa<br>cgaaaggctcagtcgaaagactgggcctttcgttttatctgttgtttgtcggtgaacgctctcctgag<br>taggacaaat |
| TetR cassette | TetR protein and Ptet promoter | ttaagacccactttcacatttaagttgtttttctaatccgcatatgatcaattcaaggccgaataaga<br>aggctggctctgcaccttggtgatcaaataattcgatagcttgtcgtaataatggcggcatactatca<br>gtagtaggtgtttccctttcttctttagcgacttgatgctcttgatcttccaatacgcaacctaaagt<br>aaaatgccccacagcgctgagtgcatataatgcattctctagtgaaaaaccttgttggcataaaaagg<br>ctaattgattttcgagagtttcatactgtttttctgtaggccgtgtacctaaatgtacttttgctcca<br>tcgcgatgacttagtaaagcacatctaaaacttttagcgttattacgtaaaaaatcttgccagctttc<br>cccttctaaagggcaaaagtgagtatggtgcctatctaacatctcaatggctaaggcgtcgagcaaag<br>cccgcttattttttacatgccaatacaatgtaggctgctctacacctagcttctgggcgagtttacgg<br>gttgttaaaccttcgattccgacctcattaagcagctctaatgcgctgttaatcactttacttttatc<br>taatctagacatcattaattcctaatttttgttgacactctatcgttgatagagttattttaccactc<br>cctatcagtgatagagaaaagaattcaaaagatctaaagaggagaaaggatct |
| PhlF cassette | PhlF protein + rrnB T1 and T7Te terminators and PphlF promoter | ctatggactatgtttgaaagggagaaatactagatggcacgtaccccgagccgtagcagcattggtag<br>cctgcgtagtccgcatacccataaagcaattctgaccagcaccattgaaatcctgaaagaatgtggtt<br>atagcggtctgagcattgaaagcgttgcacgtcgtgccggtgcaagcaaaccgaccatttatcgttgg<br>tggaccaataaagcagcactgattgccgaagtgtatgaaaatgaaagcgaacaggtgcgtaaatttcc<br>ggatctgggtagctttaaagccgatctggattttctgctgcgtaatctgtggaaagtttggcgtgaaa<br>ccatttgtggtgaagcatttcgttgtgttattgcagaagcacagctggaccctgcaaccctgacccag<br>ctgaaagatcagtttatggaacgtcgtcgtgagatgccgaaaaaactggttgaaaatgccattagcaa<br>tggtgaactgccgaaagataccaatcgtgaactgctgctggatatgattttttggtttttgttggtatc<br>gcctgctgaccgaacagctgaccgttgaacaggatattgaagaatttaccttcctgctgattaatggt<br>gtttgtccgggtacacagcgttaactagggcccataccccccaggcatcaaataaaacgaaaggctcag<br>tcgaaagactgggcctttcgttttatctgttgtttgtcggtgaacgctctctactagagtcacactgg<br>ctcaccttcgggtgggcctttctgcgtttatatacgcgacgtacggtggaatctgattcgttaccaat<br>tgacatgatacgaaacgtaccgtatcgttaaggtctagggcccataccccccaggcat |
| ***dCas9 Parts*** | | |
| PA1 | promoter [2] | tttacaccTAGCTCAGTCCTAggtattatgctagc |
| PA2 | promoter [2] | tttacaccAACGGGTCACACGggtattatgctagc |
| PA4 | promoter [2] | tttacaccTCCACAACTAGCTggtattatgctagc |
| A1NT | sgRNA [2] | ataatacctaggactgagctgttttagagctagaaatagcaagttaaaataaggctagtccgattgca<br>acttgaaaaagtggcaccgagtcggtgcttttttt |
| A2NT | sgRNA [2] | ataatacccgtgtgacccgtgttttagagctagaaatagcaagttaaaataaggctagtccgttatca<br>acttgaaaaagtggcaccgagtcggtgcttttttt |
| A4NT | sgRNA [2] | ataataccagctagttgtgggttttagagctagaaatagcaagttaaaataaggctagtccgttatca<br>acttgaaaaagtggcaccgagtcggtgcttttttt |
| L3S2P11 | terminator [11] for A1NT sgRNA, TA2 crRNA | ctcggtaccaaattccagaaaagagacgctttcgagcgtctttttcgttttggtcc |
| L3S2P55 | terminator [11] for A2NT sgRNA, TA4 crRNA | ctcggtaccaaagacgaacaataagacgctgaaaagcgtctttttcgttttggtcc |
| L3S2P21 | terminator [11] for A4NT sgRNA, TA1 crRNA | ctcggtaccaaattccagaaaagaggcctcccgaaagggggcctttttcgttttggtcc |

| | | |
|---|---|---|
| *S. pyogenes* dCas9 | RBS + dCas9 (D10A, H840A) [1] + rrnB T1 terminator | aaagaggagaaaggatctatggataagaaatactcaataggcttagctatcggcacaaatagcgtcgg<br>atgggcggtgatcactgatgaatataaggttccgtctaaaaagttcaaggttctgggaaatacagacc<br>gccacagtatcaaaaaaaatcttataggggctcttttatttgacagtggagagacagcggaagcgact<br>cgtctcaaacggacagctcgtagaaggtatacacgtcggaagaatcgtatttgttatctacaggagat<br>tttttcaaatgagatggcgaaagtagatgatagtttctttcatcgacttgaagagtcttttttggtgg<br>aagaagacaagaagcatgaacgtcatcctatttttggaaatatagtagatgaagttgcttatcatgag<br>aaatatccaactatctatcatctgcgaaaaaaattggtagattctactgataaagcggatttgcgctt<br>aatctatttggccttagcgcatatgattaagtttcgtggtcattttttgattgagggagatttaaatc<br>ctgataatagtgatgtggacaaactatttatccagttggtacaaacctacaatcaattatttgaagaa<br>aaccctattaacgcaagtggagtagatgctaaagcgattctttctgcacgattgagtaaatcaagacg<br>attagaaaatctcattgctcagctccccggtgagaagaaaaatggcttatttgggaatctcattgctt<br>tgtcattgggtttgacccctaattttaaatcaaattttgatttggcagaagatgctaaattacagctt<br>tcaaaagatacttacgatgatgatttagataatttattggcgcaaattggagatcaatatgctgattt<br>gtttttggcagctaagaatttatcagatgctattttactttcagatatcctaagagtaaatactgaaa<br>taactaaggctcccctatcagcttcaatgattaaacgctacgatgaacatcatcaagacttgactctt<br>ttaaaagctttagttcgacaacaacttccagaaaagtataaagaaatcttttttgatcaatcaaaaaa<br>cggatatgcaggttatattgatgggggagctagccaagaagaattttataaatttatcaaaccaattt<br>tagaaaaaatggatggtactgaggaattattggtgaaactaaatcgtgaagatttgctgcgcaagcaa<br>cggacctttgacaacggctctattccccatcaaattcacttgggtgagctgcatgctattttgagaag<br>acaagaagacttttatccatttttaaaagacaatcgtgagaagattgaaaaaatcttgacttttcgaa<br>ttccttattatgttggtccattggcgcgtggcaatagtcgttttgcatggatgactcggaagtctgaa<br>gaaacaattaccccatggaattttgaagaagttgtcgataaaggtgcttcagctcaatcatttattga<br>acgcatgacaaactttgataaaaatcttccaaatgaaaaagtactaccaaaacatagtttgctttatg<br>agtattttacggtttataacgaattgacaaaggtcaaatatgttactgaaggaatgcgaaaaccagca<br>tttctttcaggtgaacagaagaaagccattgttgatttactcttcaaaacaaatcgaaaagtaaccgt<br>taagcaattaaaagaagattatttcaaaaaaatagaatgttttgatagtgttgaaatttcaggagttg<br>aagatagatttaatgcttcattaggtacctaccatgatttgctaaaaattattaaagataaagatttt<br>ttggataatgaagaaaatgaagatatcttagaggatattgttttaacattgaccttatttgaagatag<br>ggagatgattgaggaaagacttaaaacatatgctcacctctttgatgataaggtgatgaaacagctta<br>aacgtcgccgttatactggttggggacgtttgtctcgaaaattgattaatggtattagggataagcaa<br>tctggcaaaacaatattagattttttgaaatcagatggttttgccaatcgcaattttatgcagctgat<br>ccatgatgatagtttgacatttaaagaagacattcaaaaagcacaagtgtctggacaaggcgatagtt<br>tacatgaacatattgcaaatttagctggtagccctgctattaaaaaaggtattttacagactgtaaaa<br>gttgttgatgaattggtcaaagtaatggggcggcataagccagaaaatatcgttattgaaatggcacg<br>tgaaaatcagacaactcaaaagggccagaaaaattcgcgagagcgtatgaaacgaatcgaagaaggta<br>tcaaagaattaggaagtcagattcttaaagagcatcctgttgaaaatactcaattgcaaaatgaaaag<br>ctctatctctattatctccaaaatggaagagacatgtatgtggaccaagaattagatattaatcgttt<br>aagtgattatgatgtcgatgccattgttccacaaagtttccttaaagacgattcaatagacaataagg<br>tcttaacgcgttctgataaaaatcgtggtaaatcggataacgttccaagtgaagaagtagtcaaaaag<br>atgaaaaactattggagacaacttctaaacgccaagttaatcactcaacgtaagtttgataatttaac<br>gaaagctgaacgtggaggtttgagtgaacttgataaagctggttttatcaaacgccaattggttgaaa<br>ctcgccaaatcactaagcatgtggcacaaattttggatagtcgcatgaatactaaatacgatgaaaat<br>gataaacttattcgagaggttaaagtgattaccttaaaatctaaattagtttctgacttccgaaaaga<br>tttccaattctataaagtacgtgagattaacaattaccatcatgcccatgatgcgtatctaaatgccg<br>tcgttggaactgctttgattaagaaatatccaaaacttgaatcggagtttgtctatggtgattataaa<br>gtttatgatgttcgtaaaatgattgctaagtctgagcaagaaataggcaaagcaaccgcaaaatattt<br>cttttactctaatatcatgaacttcttcaaaacagaaattacacttgcaaatggagagattcgcaaac<br>gccctctaatcgaaactaatggggaaactggagaaattgtctgggataaagggcgagattttgccaca<br>gtgcgcaaagtattgtccatgccccaagtcaatattgtcaagaaaacagaagtacagacaggcggatt<br>ctccaaggagtcaattttaccaaaaagaaattcggacaagcttattgctcgtaaaaaagactgggatc<br>caaaaaaatatggtggttttgatagtccaacggtagcttattcagtcctagtggttgctaaggtggaa<br>aaagggaaatcgaagaagttaaaatccgttaaagagttactagggatcacaattatggaaagaagttc<br>ctttgaaaaaaatccgattgactttttagaagctaaaggatataaggaagttaaaaaagacttaatca<br>ttaaactacctaaatatagtctttttgagttagaaaacggtcgtaaacggatgctggctagtgccgga<br>gaattacaaaaaggaaatgagctggctctgccaagcaaatatgtgaattttttatatttagctagtca<br>ttatgaaaagttgaagggtagtccagaagataacgaacaaaaacaattgtttgtggagcagcataagc<br>attatttagatgagattattgagcaaatcagtgaattttctaagcgtgttattttagcagatgccaat<br>ttagataaagttcttagtgcatataacaaacatagagacaaaccaatacgtgaacaagcagaaaatat<br>tattcatttatttacgttgacgaatcttggagctcccgctgcttttaaatattttgatacaacaattg<br>atcgtaaacgatatacgtctacaaaagaagttttagatgccactcttatccatcaatccatcactggt<br>ctttatgaaacacgcattgatttgagtcagctaggaggtgactaactcgagtaaggatctccaggcat<br>caaataaaacgaaaggctcagtcgaaagactgggcctttcgttttatctgttgtttgtcggtgaacgc<br>tctc |
| ***dCas12a Parts*** | | |
| TA1 crRNA | crRNA | TAATTTCTACTGTTGTAGATcaccTAGCTCAGTCCTAggt |
| TA2 crRNA | crRNA | TAATTTCTACTGTTGTAGATcaccAACGGGTCACACGggt |
| TA4 crRNA | crRNA | TAATTTCTACTGTTGTAGATcaccTCCACAACTAGCTggt |
| SarJ ribozyme | ribozyme spacer [12] | AGACTGTCGCCGGATGTGTATCCGACCTGACGATGGCCCAAAAGGGCCGAAACAGTCCTCTACAAATA<br>ATTTTGTTTAA |

| | | |
|---|---|---|
| PlmJ ribozyme | ribozyme spacer [12] | AGTCATAAGTCTGGGCTAAGCCCACTGATGAGTCGCTGAAATGCGACGAAACTTATGACCTCTACAAA<br>TAATTTTGTTTAA |
| *F. novicida* dCas12a | RBS + dCas12a (D917A) [3] + ECK120029600 terminator [11] | aaagaggagaaaggatctatgtcaatttatcaagaatttgttaataaatatagtttaagtaaaactct<br>aagatttgagttaatcccacagggtaaaacacttgaaaacataaaagcaagaggtttgattttagatg<br>atgagaaaagagctaaagactacaaaaaggctaaacaaataattgataaatatcatcagtttttata<br>gaggagatattaagttcggtttgtattagcgaagatttattacaaaactattctgatgtttattttaa<br>acttaaaaagagtgatgatgataatctacaaaaagattttaaaagtgcaaaagatacgataaagaaac<br>aaatatctgaatatataaaggactcagagaaatttaagaatttgtttaatcaaaaccttatcgatgct<br>aaaaaagggcaagagtcagatttaattctatggctaaagcaatctaaggataatggtatagaactatt<br>taaagccaatagtgatatcacagatatagatgaggcgttagaaataatcaaatcttttaaaggttgga<br>caacttattttaagggttttcatgaaaatagaaaaaatgtttatagtagcaatgatattcctacatct<br>attatttataggatagtagatgataatttgcctaaatttctagaaaataaagctaagtatgagagttt<br>aaaagacaaagctccagaagctataaactatgaacaaattaaaaaagatttggcagaagagctaacct<br>ttgatattgactacaaaacatctgaagttaatcaaagagtttttcacttgatgaagtttttgagata<br>gcaaactttaataattatctaaatcaaagtggtattactaaatttaatactattattggtggtaaatt<br>tgtaaatggtgaaaatacaaagagaaaaggtataaatgaatatataaatctatactcacagcaaataa<br>atgataaaacactcaaaaaatataaaatgagtgttttatttaagcaaattttaagtgatacagaatct<br>aaatcttttgtaattgataagttagaagatgatagtgatgtagttacaacgatgcaaagtttttatga<br>gcaaatagcagcttttaaaacagtagaagaaaaatctattaaagaaacactatctttattatttgatg<br>atttaaaagctcaaaaacttgatttgagtaaaatttattttaaaaatgataaatctcttactgatcta<br>tcacaacaagtttttgatgattatagtgttattggtacagcggtactagaatatataactcaacaaat<br>agcacctaaaaatcttgataaccctagtaagaaagagcaagaattaatagccaaaaaaactgaaaaag<br>caaaatacttatctctagaaactataaagcttgccttagaagaatttaataagcatagagatatagat<br>aaacagtgtaggtttgaagaaatacttgcaaactttgcggctattccgatgatatttgatgaaatagc<br>tcaaaacaaagacaatttggcacagatatctatcaaatatcaaaatcaaggtaaaaaagacctacttc<br>aagctagtgcggaagatgatgttaaagctatcaaggatcttttagatcaaactaataatctcttacat<br>aaactaaaaatatttcatattagtcagtcagaagataaggcaaatattttagacaaggatgagcattt<br>ttatctagtatttgaggagtgctactttgagctagcgaatatagtgcctctttataacaaaattagaa<br>actatataactcaaaagccatatagtgatgagaaatttaagctcaattttgagaactcgactttggct<br>aatggttgggataaaaataaagagcctgacaatacggcaattttatttatcaaagatgataaatatta<br>tctgggtgtgatgaataagaaaaataacaaaatatttgatgataaagctatcaaagaaaataaaggcg<br>agggttataaaaaaattgtttataaacttttacctggcgcaaataaaatgttacctaaggttttcttt<br>tctgctaaatctataaaattttataatcctagtgaagatatacttagaataagaaatcattccacaca<br>tacaaaaaatggtagtcctcaaaaaggatatgaaaaatttgagtttaatattgaagattgccgaaaat<br>ttatagatttttataaacagtctataagtaagcatccggagtggaaagattttggatttagattttct<br>gatactcaaagatataattctatagatgaattttatagagaagttgaaaatcaaggctacaaactaac<br>ttttgaaaatatatcagagagctatattgatagcgtagttaatcagggtaaattgtacctattccaaa<br>tctataataaagatttttcagcttatagcaaagggcgaccaaatctacatactttatattggaaagcg<br>ctgtttgatgagagaaatcttcaagatgtggtttataagctaaatggtgaggcagagctttttatcg<br>taaacaatcaatacctaaaaaatcactcacccagctaaagaggcaatagctaataaaaacaaagata<br>atcctaaaaagagagtgtttttgaatatgatttaatcaaagataaacgctttactgaagataagttt<br>ttctttcactgtcctattacaatcaattttaaatctagtggagctaataagtttaatgatgaaatcaa<br>tttattgctaaaagaaaaagcaaatgatgttcatatattaagtatagCtagaggtgaaagacatttag<br>cttactatactttggtagatggtaaaggcaatatcatcaaacaagatactttcaacatcattggtaat<br>gatagaatgaaaacaaactaccatgataagcttgctgcaatagagaaagatagggattcagctaggaa<br>agactggaaaaagataaataacatcaaagagatgaaagagggctatctatctcaggtagttcatgaaa<br>tagctaagctagttatagagtataatgctattgtggtttttgaggatttaaattttggatttaaaaga<br>gggcgtttcaaggtagagaagcaggtctatcaaaagttagaaaaaatgctaattgagaaactaaacta<br>tctagttttcaaagataatgagtttgataaaactgggggagtgcttagagcttatcagctaacagcac<br>cttttgagacttttaaaaagatgggtaaacaaacaggtattatctactatgtaccagctggttttact<br>tcaaaaatttgtcctgtaactggttttgtaaatcagttatatcctaagtatgaaagtgtcagcaaatc<br>tcaagagttctttagtaagtttgacaagatttgttataaccttgataagggctattttgagtttagtt<br>ttgattataaaaactttggtgacaaggctgccaaaggcaagtggactatagctagctttgggagtaga<br>ttgattaactttagaaattcagataaaaatcataattgggatactcgagaagtttatccaactaaaga<br>gttggagaaattgctaaaagattattctatcgaatatgggcatggcgaatgtatcaaagcagctattt<br>gcggtgagagcgacaaaaagtttttgctaagctaactagtgtcctaaatactatcttacaaatgcgt<br>aactcaaaaacaggtactgagttagattatctaatttcaccagtagcagatgtaaatggcaatttctt<br>tgattcgcgacaggcgccaaaaaatatgcctcaagatgctgatgccaatggtgcttatcatattgggc<br>taaaaggtctgatgctactaggtaggatcaaaaataatcaagagggcaaaaaactcaatttggttatc<br>aaaaatgaagagtattttgagttcgtgcagaataggaataactaattgagaagagaaaagaaaaccgc<br>cgatcctgtccaccgcattactgcaaggtagtggacaagaccggcggtcttaagtttttggctgaa |

(red: ribozyme, orange: gene, purple: terminator, green: promoter)

**Table S5. Primers.** Select primers used for cloning.

| Use | Name | DNA Sequence (5' to 3') |
|---|---|---|
| Reporters | dCR-PA4-F | atat actagtatgcgtctcctttacacc tccacaactagct ggtattatgctagcagctg |
| | dCR-PA1-F | atat actagtatgcgtctcctttacacc TAGCTCAGTCCTA ggtattatgctagcagctg |
| | dCR-PA2-F | atat actagtatgcgtctcctttacacc AACGGGTCACACG ggtattatgctagcagctg |
| | Vs-RiboJ-R | TGCGGTACCTTTCTCCTCTTTGTAAAGTTAAACAAAATTATTTGTAGAGGCTGTTTCGTC |
| | dCR-Venus-F | CTTTACAAAGAGGAGAAAGGTACCGC |
| | XbaI-rrnB-R | TTAATCTAGAATTTGTCCTACTCAGGAGAGCG |
| sgRNAs | 385-BglII-F | ttaa agatctcagaaatcattttacaccaacgg |
| | 385-HindIII-R | atat AAGCTTAAGCGGACCAAAACGAAAAAAGG |
| | 386-BgH-F | ttaa AGATCTAAGCTTTTTACACCTCCACAACTAGCTGG |
| | 386-Bam-R | atat GGATCCTATCGGACCAAAACGAAAAAAG |
| | 387-BB-PA1-F | ttaa AGATCTGGATCCTTTACACCTAGCTCAGTCCTAGGTATTATGCTAGCATAATACCCGTGTGACCCGT |
| | 387-Xba-R | atat TCTAGATAAAGGACCAAAACGAAAAAAGACGC |
| pJK398 | 398-388-R | gagtcagtgagcgaggaagcctgcagtctagataa |
| | 398-p15A-F | gactgcaggcttcctcgctcactgactcgctac |
| | 398-p15A-R | cgtgagttttcgttccactgccaagcactagtaacaactt |
| | 398-388-F | gttactagtgcttggcagtggaacgaaaactcacg |
| PphlF sgRNA (pJK410, 411) | 410-Bg-PhlF-F | TTAA AGATCT ACTTTTACAGCTAGCTCAGTCC |
| | 410-Hind-R | ATAT AAGCTT CTTAAGTTTTTTGGCTGAAGGACC |
| | 411-Bam-PhlF-F | TTAA ggatcc ACTTTTACAGCTAGCTCAGTCC |
| | 411-Pst-R | ATAT CTGCAG GAATTCTAAAGGACCAAAACGA |
| ssRA-tagged dCas9 | dCas9-DENYA-R | AGCGTAGTTTTCGTCGTTTGCTGCGTCACCTCCTAGCTGACTCAAATCAATGCG |
| | LAA-F | GCAGCAAACGACGAAAACTACGCT TTAGCAGCT TAACTCGAGTAAGGATCTCCAGGCATC |
| | LVA-F | GCAGCAAACGACGAAAACTACGCT TTAGTAGCT TAACTCGAGTAAGGATCTCCAGGCATC |
| | ASV-F | GCAGCAAACGACGAAAACTACGCT GCATCAGTT TAACTCGAGTAAGGATCTCCAGGCATC |
| | AAV-F | GCAGCAAACGACGAAAACTACGCT GCAGCAGTT TAACTCGAGTAAGGATCTCCAGGCATC |
| | SC101-R | ACACCAGAACAGCCCGTTTG |
| | 414-chk-F | TAGGAGGTGACGCAGCAAAC |
| | 414-chk-Rv | AAGCTGTTCACCATGAACAG |
| dCRv1 clonetegration (pJK422 + 423) | SpeI-398-F | ATATATactagtaataggcgtatcacgaggcaga |
| | Bss-412-F | TATAGCGCGCCTAGTATGCGTCTCCTTTACACC |
| | SpeI-412-R | AATTACTAGTAGCAAAACCCGTACCCTAGAT |
| | SpeI-413-Rv | AATTACTAGTAATAGGCGTATCACGAGGCA |
| | SpeI-It0Vs-R | AATTACTAGTGACTCCTGTTGATAGATCCAGTA |
| | Bss-425-412-F | TATAGCGCGCTACTAGAGTCACACTGGCTCAC |
| | SpeI-427-R | ATATATACTAGTATAAACGCAGAAAGGCCCACC |
| | Bss-428-F | TATAGCGCGCACGTCTCATTTTCGCCAGATATCG |
| | 433end-R | ATCTTCCAGGAAATCTCCGC |
| | SC101R2 | GCGATGCTGCTTATCGA |
| | 435-SpeI-R | AATTACTAGTAGCAAAACCCGTACCCTAGA |
| sgRNA Set 2+3 (1,3,5) | 440-PA5-F | ATAT AGATCTCAGAAATCATTTTACACCAAAACACTCGGAGGGTATTATGCTAGCA |
| | 445-PA5-F | TACACCAAAACACTCGGAGGGTATTATGCTAGCAGCTG |
| | 445-PA5-R | TAATACCCTCCGAGTGTTTTGGTGTAAAGGAGACGCATAC |
| | 446-PA3-F | TACACCCGAAATGGAGCATGGTATTATGCTAGCAGCTG |
| | 446-PA3-R | TAATACCATGCTCCATTTCGGGTGTAAAGGAGACGCATAC |
| dCas12a | 500-vR | ataaattgacatagatcctttctcctcttt |
| | 500-Cp-F | ggatctatgtcaatttatcaagaatttg |
| | 500-Cp-R | gctaaatgtctttcacctctaGctatacttaatatatgaacatcatttgc |
| | 500-Cp-2F | atagCtagaggtgaaagacatttagc |

| | | |
|---|---|---|
| | 500-Cp-2R | tagttattcctattctgcacgaactc |
| | 500-vF | TTAATTAAgtacgggttttgctgcccg |
| | 504-v-F | CCGTTAATTAACTGCAGGCTTCCTCGCTCACTGACTCGCTAC |
| | 504-v-R | CCGTTAATTAAATGGTTTCTTAGACGTCAGG |
| | 516-105-F | tttacggctagctcagtcctaggtactatgctagcagtacgtctg |
| | 516-Phlf-R | AggtgATCTACAACAGTAGAAATTAaccttaacgatacggtacg |
| | 516-v-F | TCTACTGTTGTAGATCACCTAGCTCAGTCCTAGGTCTCGG |
| | 516-v-R | TACCTAGGACTGAGCTAGCCGTAAAATGATTTCTGAGATC |
| | 517-v-F | TCTACTGTTGTAGATcaccTCCACAACTAGCTggtctcgg |
| | 517-v-R | tacctaggactgagctagccgtaaaatcctatcTTAAACA |
| | 528-511-F | cagtatgcgtatttgcgcgcttggcctagtatgcgtctcc |
| | 528-v-R | ggagacgcatactaggccaagcgcgcaaatacgcatactg |
| | 528-511-R | ccatgcatctcgaggcatgcacaccagaacagcccgtttg |
| | 528-v-F | caaacgggctgttctggtgtgcatgcctcgagatgcatgg |
| PhlF-TA1 | 520-phlF-F | ATCTACAACAGTAGAAATTAACCTTAACGATACGGTACGTT |
| | 520-phlF-R | TGATTTCTGGAATTCTAAAGATCTTTTACGGCTAGCTCAGTC |
| | 520-v-F | AGATCTTTAGAATTCCAGAAATCATCCTTAGC |
| | 520-v-R | GTTAATTTCTACTGTTGTAGATCACCTAGCTCAGTCCTAGGT |

#### References

(1) Qi, L. S.; Larson, M. H.; Gilbert, L. A.; Doudna, J. A.; Weissman, J. S.; Arkin, A. P.; Lim, W. A. Repurposing CRISPR as an RNA-guided platform for sequence-specific control of gene expression. *Cell* **2013**, *152*, 1173.

(2) Nielsen, A. A.; Voigt, C. A. Multi-input CRISPR/Cas genetic circuits that interface host regulatory networks. *Mol Syst Biol* **2014**, *10*, 763.

(3) Zetsche, B.; Gootenberg, J. S.; Abudayyeh, O. O.; Slaymaker, I. M.; Makarova, K. S.; Essletzbichler, P.; Volz, S. E.; Joung, J.; van der Oost, J.; Regev, A.; Koonin, E. V.; Zhang, F. Cpf1 is a single RNA-guided endonuclease of a class 2 CRISPR-Cas system. *Cell* **2015**, *163*, 759.

(4) Gibson, D. G.; Young, L.; Chuang, R. Y.; Venter, J. C.; Hutchison, C. A.; Smith, H. O. Enzymatic assembly of DNA molecules up to several hundred kilobases. *Nat Methods* **2009**, *6*, 343.

(5) St-Pierre, F.; Cui, L.; Priest, D. G.; Endy, D.; Dodd, I. B.; Shearwin, K. E. One-step cloning and chromosomal integration of DNA. *ACS Synth Biol* **2013**, *2*, 537.

(6) Luro, S.; Potvin-Trottier, L.; Okumus, B.; Paulsson, J. Isolating live cells after high-throughput, long-term, time-lapse microscopy. *Nat Methods* **2020**, *17*, 93.

(7) Potvin-Trottier, L.; Lord, N. D.; Vinnicombe, G.; Paulsson, J. Synchronous long-term oscillations in a synthetic gene circuit. *Nature* **2016**, *538*, 514.

(8) Skinner, S. O.; Sepulveda, L. A.; Xu, H.; Golding, I. Measuring mRNA copy number in individual Escherichia coli cells using single-molecule fluorescent in situ hybridization. *Nat Protoc* **2013**, *8*, 1100.

(9) Jones, D. L.; Leroy, P.; Unoson, C.; Fange, D.; Curic, V.; Lawson, M. J.; Elf, J. Kinetics of dCas9 target search in Escherichia coli. *Science* **2017**, *357*, 1420.

(10) Andersen, J. B.; Sternberg, C.; Poulsen, L. K.; Bjorn, S. P.; Givskov, M.; Molin, S. New unstable variants of green fluorescent protein for studies of transient gene expression in bacteria. *Appl Environ Microbiol* **1998**, *64*, 2240.

(11) Chen, Y. J.; Liu, P.; Nielsen, A. A.; Brophy, J. A.; Clancy, K.; Peterson, T.; Voigt, C. A. Characterization of 582 natural and synthetic terminators and quantification of their design constraints. *Nat Methods* **2013**, *10*, 659.

(12) Lou, C.; Stanton, B.; Chen, Y. J.; Munsky, B.; Voigt, C. A. Ribozyme-based insulator parts buffer synthetic circuits from genetic context. *Nat Biotechnol* **2012**, *30*, 1137.
